# Turning an object into a scene: buildings activate scene-selective visual cortex independently of visual features

**DOI:** 10.64898/2026.07.12.738103

**Authors:** Yuanfang Zhao, Simen Hagen, Marius V. Peelen

**Affiliations:** Donders Institute for Brain, Cognition and Behaviour, Radboud University, Nijmegen, The Netherlands; Department of Cognitive Science, Johns Hopkins University, Baltimore, USA

**Keywords:** Object representation, Scene perception, Parahippocampal place area, Ultra-fast fMRI, Category selectivity

## Abstract

Human visual cortex contains regions that selectively respond to both scenes and large objects, particularly buildings. The cortical overlap between buildings and scenes has been attributed to shared visual features (e.g., cardinal orientations, rectilinearity). Alternative accounts propose that buildings may also activate scene representations indirectly, independently of specific visual features, for example because buildings evoke a sense of space. Here, we tested for such feature-independent activation by comparing EEG and fMRI responses in human participants (both sexes) to buildings and visually-matched boxes, and relating these responses to scene-selective responses. Buildings and boxes were matched using image-based metrics, deep neural networks, and a perceptual similarity task. Time-resolved EEG decoding showed that buildings and boxes evoked discriminable responses from 360ms post-stimulus onset, incompatible with feedforward visual feature processing. Importantly, the building-box classifier generalized to discriminate scenes from chairs, providing the first EEG evidence for a representational overlap between buildings and scenes. Temporal generalization analyses further showed that the late building-selective response corresponded to an earlier scene-selective response, with a temporal offset of ∼130ms. Finally, ultra-fast fMRI (TR=140ms) revealed that the scene-selective parahippocampal place area (PPA) similarly showed a feature-independent building-selective response that was delayed and prolonged relative to the scene-selective response. These results clarify the nature of building selectivity in visual cortex by showing that this selectivity is feature-independent and delayed relative to scene selectivity. We propose that intrinsic associations between object categories (e.g., buildings and scenes, tools and hands) may account for part of the categorical organization of ventral temporal cortex.

**Significance Statement:** The human ventral stream contains regions responding selectively to behaviorally-relevant categories such as places, faces, and words. Contrasting views attribute this selectivity either to the feedforward processing of visual features or to top-down input from domain-specific networks. Here, we show that neural selectivity for buildings persists in the absence of category-diagnostic visual features. EEG decoding showed that this high-level building response emerged late during object processing and corresponded to an earlier scene-selective response. Ultra-fast fMRI revealed that these findings were also observed in the scene-selective parahippocampal place area (PPA). These results show that selectivity for some categories can be decoupled from visual feature processing, reflecting associative processes that follow object recognition.

## Introduction

Human visual cortex shows a categorical organization, with spatio-temporal response patterns evoked by pictures of objects organized along dimensions of animacy and real-world size (Kriegeskorte et al., 2008; Konkle and Oliva, 2012; Carlson et al., 2013), and with focal selectivity for more specific categories of stimuli, including scenes, faces, bodies, words, and tools (Downing et al., 2006). This organization has been widely replicated using multiple methods, including fMRI and M/EEG, but the driving factors are still debated (Haxby et al., 2001; Op de Beeck et al., 2008; Peelen and Downing, 2017; Ritchie et al., 2026).

One of the first reports of category selectivity using fMRI was for buildings (Aguirre et al., 1998), in a region that overlaps scene-selective cortex (Epstein and Kanwisher, 1998). More recently, studies have shown that this selectivity extends to large/stable objects more generally (e.g., furniture; Mullally and Maguire, 2011; Konkle and Oliva, 2012; He et al., 2013; Troiani et al., 2014; Hagen et al., 2026). Here, using EEG and ultra-fast fMRI, we aimed to shed light on the temporal dynamics and representational overlap between building and scene selectivity.

One possibility is that the selective response to buildings reflects the feedforward processing of visual features typical of such objects (e.g., rectilinear features, cardinal orientations), which may be shared with scenes (Nasr and Tootell, 2012; Nasr et al., 2014). For example, fMRI studies have shown that large-object-selective responses remain when stimuli are made unrecognizable while preserving low- and mid-level features, suggesting that this selectivity reflects the feedforward processing of such features (Long et al., 2018). These findings were confirmed by studies using time-sensitive M/EEG decoding, showing that large and small objects evoke distinct response pattern as early as 150 ms after stimulus onset (Khaligh-Razavi et al., 2018), even when made unrecognizable (Wang et al., 2022). This early time course suggests that large-object selectivity at least partly reflects visual feature processing.

Other studies, however, have observed selectivity for buildings and large objects when participants merely imagined such objects (Ishai et al., 2000; Mullally and Maguire, 2011; Boccia et al., 2021), raising the possibility that this selectivity could reflect a relatively abstract representation of space, context, or navigation-related cognitive processes. Specifically, Mullally and Maguire (2011) proposed that buildings and large/stable objects evoke a sense of surrounding space and that neural selectivity to buildings reflects this sense of surrounding space (see also Bainbridge & Oliva, 2015 for the related concept of interaction envelope). Such an account naturally explains the overlap between building and scene selectivity but would predict a delayed response to buildings relative to scenes.

In the present study, we aimed to shed light on the nature of building selectivity, its relation to scene selectivity, and their temporal dynamics. To do so, we developed a carefully controlled stimulus set consisting of buildings and visually-matched controls (boxes), and used EEG and ultra-fast fMRI to test: 1) whether neural selectivity to buildings can be observed when buildings are contrasted with visually-matched non-building objects (boxes); 2) whether these building-selective responses correspond to scene-selective responses; and 3) whether the building-selective responses are delayed relative to scene-selective responses. As a preview of the results, in both EEG and fMRI experiments, we observed a clear building-selective response after equating perceptual similarity and controlling for low- and mid-level visual features across categories. In EEG, this response emerged relatively late, from around 360 ms after stimulus onset, ruling out that it was driven by the feedforward processing of visual features and instead suggesting a top-down origin. Furthermore, results revealed that the building-selective response pattern corresponded to the scene-selective response pattern, with a temporal offset of ∼130 ms, suggesting that buildings indirectly activated scene representations. Finally, fMRI showed a building-selective response in the scene-selective PPA that was delayed relative to the scene-selective response. We interpret these results as reflecting the automatic association of buildings and scenes, such that seeing a building primes the representation of scenes (or space).

## Methods

### Behavioral study

The behavioral study used an oddball visual search task (Arun, 2012) that provides a sensitive measure of the pairwise perceptual similarity of the objects that were included in the neuroimaging experiments.

#### Participants

A total of 45 participants were recruited via Prolific (www.prolific.com) in exchange for monetary compensation. Five participants were rejected due to not being in full-screen mode for at least 10% of the trials while conducting the experiment, leaving 40 participants for the analysis (20 females, mean age = 27 ± 3.7 years). All participants reported having normal or corrected-to-normal vision and no history of neurological or psychiatric disorders. All participants provided digital informed consent before the experiment. The experiment was approved by Radboud University Faculty of Social Sciences Ethics Committee (ECSW2017–2306-517).

#### Stimuli

Stimuli consisted of 8 images of buildings and 8 images of boxes (Figure 1A). Notably, images of buildings and boxes were carefully selected so that each building exemplar was matched to a box exemplar in terms of overall contour shape and viewing angle. These images were resized to the same size and fine-tuned in Photoshop to minimize any contour shape and perspective differences. This process aimed to eliminate potential low- and mid-level visual feature differences between buildings and boxes while preserving their diagnostic visual features (e.g., windows/doors for buildings and hinged lids and locks for boxes). An independent online pilot study (N = 40) further ensured the images of buildings and boxes were highly recognizable when viewed one by one (as in the neuroimaging experiments).

**Figure 1.**
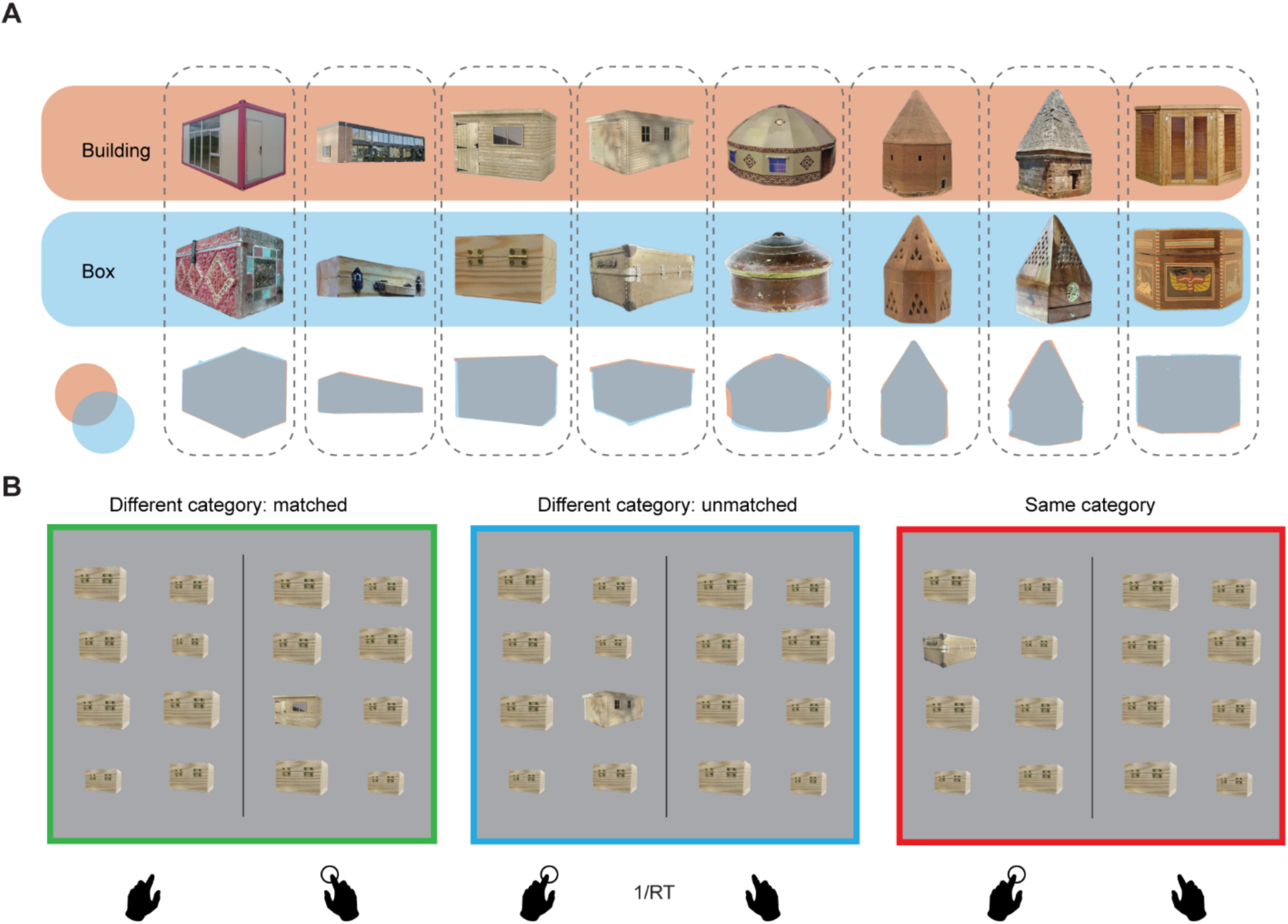
Experimental materials and examples of search arrays for the behavioral experiment. (A) Visually matched images of buildings and boxes used in behavioral, EEG, and fMRI experiments. The bottom row illustrates the close matching of the outlines across the two categories. (B) Examples of the search arrays for the behavioral experiment. Left: a search array where the oddball target is a building and the distractor is a visually matched box (matched objects); Middle, a search array where the oddball target is a building and the distractor is a box, but the box is not visually matched to that building (unmatched objects); Right, a search array where both the oddball target and the distractor are boxes (same category objects). 1/RT represents the multiplicative inverse of the reaction time (RT).

#### Experimental design

The experiment was programmed using JsPsych 6.1.0 (de Leeuw, 2015) and presented online through the hosting service provided by Pavlovia (https://pavlovia.org/, Open Science Tools, Nottingham, UK). In each trial, following an initial 700 ms fixation period, participants viewed an array of spatially arranged objects. These objects were organized into 4 rows and 2 columns on each side of the monitor, separated by a vertical straight line at the central position (Figure 1B). Each array included one “oddball” target object, positioned either to the right or left of the central line, among other “distractor” objects. The distractor objects, identical in appearance, always differed from the target object. The location of the “oddball” target within its side was randomized. Furthermore, to avoid floor effects, we randomly jittered each objects’ retinal size. The search array remained onscreen for 5 s, during which participants were instructed to indicate the location (left or right) of the “oddball” target by pressing Z (for “left”) and M (for “right”) as quickly as possible. Participants’ response ended the search array and initiated the next trial. With 8 objects in each of the two categories, any unique “target-distractor” assignment resulted in a total of 240 combinations (i.e., 16 × 15).

Each target appeared on each side once, culminating in a total of 480 trials (i.e., 240 × 2). The trials were displayed in random order and divided into 10 equal-length blocks. Note that if participants made an erroneous response in a trial, that trial (with randomized within-side location and jittered size) would be appended at the end of the initial 480 trials until we had collected all correct responses for each target-distractor combination with the target on each side. On average, participants completed the session with 489 ± 7.8 trials in about 30 minutes.

#### Data analysis

For each unique combination of two objects, we calculated the average reaction time (RT) across the two reversed “target-distractor” assignments and the left/right appearances of the target in each assignment. Following previous work (Arun, 2012), 1/RT served as the perceptual distance metric between each pair of objects. This approach allowed us to measure the perceptual distance between pairs of objects for each participant, resulting in a symmetric perceptual distance matrix. These matrices were highly consistent across participants, with a mean Spearman-Brown-corrected split-half reliability of 0.987 (95% interval: 0.984 – 0.990; 1000 random splits of participants). The matrices were then averaged across participants.

#### Statistical testing

We categorized the unique two-object combinations into three groups: visually matched different-category objects (building-box), unmatched different-category objects (building-box), and unmatched same-category objects (building-building or box-box), and then compared the perceptual distances among these groups. A non-parametric testing approach was employed, involving the shuffling of the perceptual distance matrix values 10,000 times while maintaining its symmetry and ensuring diagonal zeros. In each shuffle, the average perceptual distance across different combinations in each group was recorded. We then compared the actual average score differences between matched and unmatched different-category objects (the effect of visual similarity) and between unmatched different-category and unmatched same-category objects (the effect of categorical similarity), to the corresponding null distributions constructed from the random shuffling. P-values below 0.05 (two-tailed) were considered significant.

### EEG study

#### Participants

We aimed to include 34 participants for 80% power to detect a medium effect size (d = 0.5) or larger. A total of 36 participants were recruited through SONA, a psychology research-related participants pool based at Radboud University, in exchange for monetary compensation. However, the data from three participants were corrupted due to malfunctions in the data collection process (one due to trigger failure, another due to battery depletion mid-test, and the third due to an exceptionally high impedance level), leaving 33 participants for the preprocessing (21 females, mean age = 29 ± 8.9 years). In the preprocessing, one participant was further rejected due to excessive data noise (see the Preprocessing section below), leaving 32 participants for the final analysis. All participants reported having normal or corrected-to-normal vision and no history of neurological or psychiatric disorders. All participants provided written informed consent before the experiment. The experiment was approved by Radboud University Faculty of Social Sciences Ethics Committee (ECSW2017–2306-517).

#### Stimuli

Stimuli consisted of 56 individual colored object (and scene) images taken from 7 basic-level categories (buildings, boxes, scenes, chairs, tools, instruments, and hands). Images of tools, instruments and hands were included for another experiment with a different purpose. Each category contained 8 exemplars. Notably, images of buildings and boxes were the same images as those used in the behavioral experiment. The images of buildings and boxes were further superimposed on a phase-scrambled background to equate the high spatial frequency (high-SF) component (defined as 5 cycles/degree, as in (Rajimehr et al., 2011). Luminance was also matched for buildings and boxes using the SHINE color toolbox (Dal Ben, 2023). Examination of image-level visual property metrics (such as rectilinearity, as in Nasr et al., 2014), activations from the layers of a deep neural network (AlexNet, Krizhevsky et al., 2012), and comparison of mid-level visual features from a texture statistic model (P-S model, Portilla and Simoncelli, 2000; Henderson et al., 2023) confirmed that visual features were overall matched between buildings and boxes (Figures S1, S2). Colored images of other categories were similarly resized and cropped to maintain a consistent retinal size with buildings/boxes and superimposed on a phase-scrambled background. Scenes images included 4 man-made scenes and 4 natural scenes, specifically selected to be devoid of large objects (e.g., buildings) and cropped by an aperture with the same diameter size as images of other categories. During the experiment, the stimuli were centrally displayed on a uniform gray background. The phase-scrambled background extended approximately 20 x 20 degrees and objects (and scenes) extended approximately 8 x 8 degrees.

#### Experimental design

The experiment took place in a separate testing room, which the experimenter monitored visually via webcam from the adjacent room. We used PsychoPy to display stimuli on a PC monitor (60 Hz refresh rate). Each image was displayed one-by-one for 500 ms, followed by a fixation cross for a randomly selected duration of 1,000, 1,100, or 1,200 ms. After approximately every 6±3 images, a probe stimulus within a blue frame was displayed (Figure 2). Participants were asked to indicate whether the probe stimulus had been presented among the immediately preceding 6±3 images by pressing F (“just seen”) or J (“not just seen”). For half of the probe stimuli, the stimulus was among the previous 6±3 images, and in the other half, the stimulus was randomly drawn from the rest of the image pool minus the previously displayed 6±3 images. The probe stimulus remained until participants made a response, after which visual feedback on the accuracy of participants’ response was immediately displayed. Participants were invited to take a break after every 31 images. There was a total of 616 trials, divided into 20 blocks, with each of the 56 stimuli repeated 11 times. Trial order was random. The entire session lasted approximately 30 minutes (mean = 29.50 min, range = 24.65–36.89 min).

**Figure 2.**
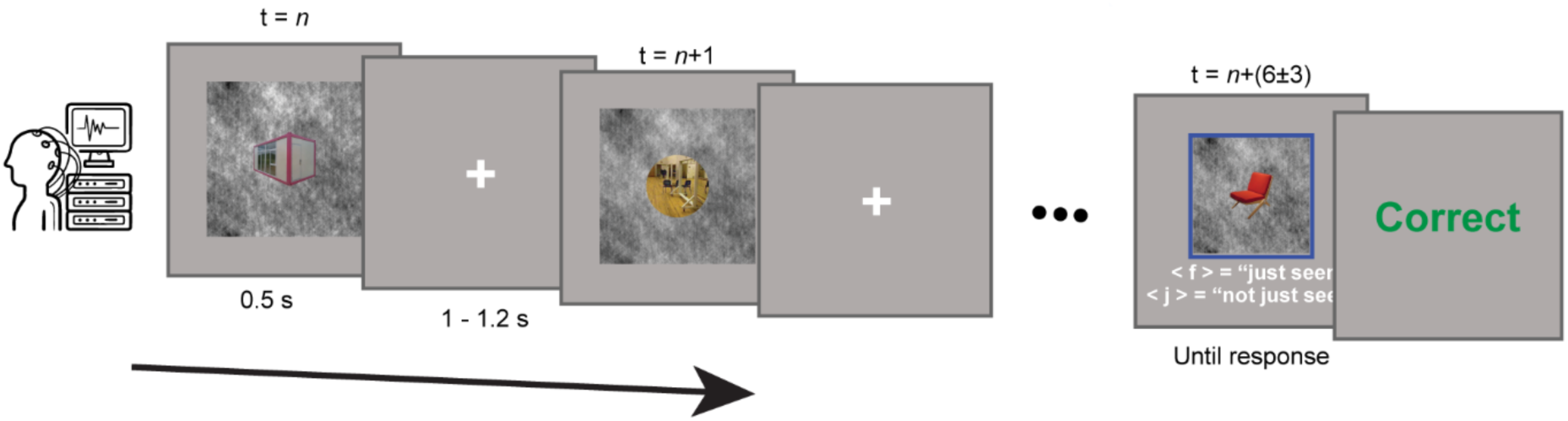
EEG experiment procedures. Schematic illustrating the EEG experiment procedure. Participants viewed objects and scenes on top of a phase-scrambled background, presented at fixation. They were asked to perform a simple memory recognition task after every 6 trials to ensure they remained attentive.

#### EEG Acquisition

We recorded scalp EEG using a 64-channel active electrode actiCAP system (500 Hz sample rate) with customized electrode positions adapted from the actiCAP 64Ch Standard-2 system (ground electrode placed at AFz; TP9 placed on the left mastoid to serve as a reference electrode; for all except one participant, to monitor eye movements, FT9/10 were placed at the outer canthi of both eyes (horizontal) and Fp1/2 were placed immediately above and below the right eye (vertical)). Electrode offsets were verified to be between 20 and 50 kOhm before recording commenced. Data were referenced online to the left mastoid and filtered between 0.016 and 125 Hz using BrainVision Recorder. We synchronized the EEG recording with the onset of each stimulus by sending markers via a USB port to the EEG acquisition PC. In the beginning of the experiment, we visually monitored the EEG trace, manually initiating each session only after it had been free from muscular and ocular artifacts for at least 5 seconds.

#### Preprocessing

We carried out EEG data preprocessing in Python 3.8 using the MNE 1.0.3 toolbox (Gramfort et al., 2013). A bipolar horizontal EOG derivation was computed as the difference between the two horizontal EOG electrodes (FT9/10), and a vertical EOG derivation was computed as the difference between the two vertical EOG electrodes (Fp1/2). For one participant without EOG channels, to maintain consistency with other participants, we removed Fp2, FT9/10 and set Fp1 as the EOG channel. For the remaining 60 electrodes, we visually identified artifact-ridden channels or non-responsive channels and replaced them using spherical spline interpolation (on average 2.9 channels/participant replaced, maximum 8 channels and minimum 0 channel). For the 60 electrodes after interpolation, we applied a finite impulse response (FIR) bandpass filter (0.05–120 Hz) to the raw EEG trace, followed by an FFT multi-notch filter to remove electrical line noise at 50 and 100 Hz. We re-referenced each channel’s signal to the average of all 60 scalp channels before segmenting each participant’s continuous EEG data for each trial from -200 to 800 ms relative to the onset of the image and resampling at 125 Hz. Epochs with deflections > 200 μV were visually examined and discarded if necessary. One participant was rejected due to an exceptionally high number of epochs removed (39, for the remaining participants, on average 2.1 trials removed, maximum 9 trials), leaving 32 participants for the analyses.

ICA was then performed on the retained epoch EEG data for each participant to visually identify and manually remove components associated with blinks and eye movements, as well as components with clearly non-neuronal origins (on average 8.9 components removed, maximum 16 and minimum 2). The ICA-denoised epoch EEG data were then baseline-corrected using the mean voltage from -200 to 0 ms relative to the onset of the image in each trial.

#### Decoding Analyses

For the subsequent statistical analyses, we focused exclusively on the 19 posterior channels of interest (P7, P5, P3, P1, Pz, P2, P4, P6, P8, PO9, PO7, PO3, POz, PO4, PO8, PO10, O1, Oz, O2). This a priori posterior channel selection was based on a large number of studies showing that neural responses in visual areas are mostly sampled by posterior channels (e.g., Romei et al., 2008). In addition, we also conducted data analysis with all channels and observed the same pattern of results (Figures S4, S5). A support vector machine classifier was trained based on neural activation patterns across posterior channels, at each time point. Standardization of the training features was implemented by removing the mean and scaling to unit variance before applying the classifier with scikit-learn using default parameters (for details: https://scikit-learn.org/stable/modules/generated/sklearn.svm.SVC.html). Decoding analyses were performed on single-trial data.

##### Leave-one-pair-out Decoding

To discriminate buildings vs. boxes, we employed exemplar-pair cross-validation, which requires the classifier to generalize to a pair of visually-matched building and box exemplars that was not included in the training set. In each fold, the trials for 7 building exemplars and their 7 visually-matched box counterparts were used to train the classifier, which was then tested on the trials from the held-out 1 building exemplar and its visually-matched box counterpart. For each fold, we measured the area under the Receiver Operating Characteristic Curve (AUC ROC), which reflects a summary measure of the model’s performance across various classification thresholds. An AUC from 0.5 to 1 represents a model with random (or chance-level) to perfect classification performance. The above procedure in a fold was iterated 8 times traversing all the 8 visually-matched building and box pairs. After completing all iterations of cross-validation, we averaged the AUC across all iterations as the final decoding performance. After this procedure was applied to each time point, the averaged decoding AUC values were smoothed across time points to minimize noise using a nine-point moving window (equivalent to a time window of ±32 ms, Bae and Luck, 2018).

To differentiate between scenes and chairs, we adopted an approach akin to that used for decoding buildings vs. boxes. Importantly, since scenes and chairs were not initially visually paired, we randomly paired exemplars from the scenes category with those from the chairs category. Subsequent procedures remained identical to those used in the buildings vs. boxes decoding. The pairings were randomized and repeated 10 times to avoid any unusual combinations. Decoding performance was then averaged across all repetitions.

##### Cross-decoding

To investigate whether activity patterns distinguishing buildings vs. boxes were related to scene processing, we trained a classifier for buildings vs. boxes using trials from all 8 pairs of visually-matched exemplars at each time point and tested it on scenes vs. chairs trials at matching time points. After this procedure was applied to each time point, the decoding AUC values were smoothed across time points to minimize noise using a nine-point moving window (equivalent to a time window of ±32 ms, Bae and Luck, 2018).

##### Temporal Generalization Analysis

To further explore the similarities between building and scene representations across time, we utilized a temporal generalization approach (King and Dehaene, 2014). Specifically, we tested the classifier trained for building vs. box using trials from all 8 pairs of visually-matched exemplars at each time point (i) on scene and chair trials, at different time points (j). If a classifier trained for building vs. box at one time point (i) can be generalized to scene vs. chair at another time point (j), it suggests that the distinguishable features are similar across building vs. box and scene vs. chair for these two specific time points (i, j). An AUC score was measured to quantify this generalizability at (i, j).

By traversing all possible time point pairs from the training set to the testing set, the resulting temporal generalization matrix may provide a more intuitive indication of this representational temporal shift. For example, above-chance generalization off the diagonal indicates an earlier or later representation in one condition compared to the other. After obtaining the temporal generalization matrix, we applied a nine-point 2D uniform filter to smooth the decoding AUC values, minimizing noise in the process.

#### Statistical Testing

We assessed whether buildings (or scenes) could be distinguished from boxes (or chairs) and whether the classifier trained for buildings vs. boxes could be generalized to scenes vs. chairs in the -200 to 800 ms interval using a non-parametric cluster-based permutation test (Maris and Oostenveld, 2007) as implemented in the MNE toolbox. This method controls the family-wise error rate (FWE) in multiple comparisons. The AUC value of every sample (time point) was compared with 0.5 (chance) using a one-sample t-statistic. Then, adjacent samples were clustered based on a t-value threshold corresponding to an uncorrected p-value of 0.05, one-tailed (given that decoding relative to chance can, in principle, only be positive). The resulting clusters’ FWE-corrected p-value was computed using the Monte Carlo method involving 10,000 random sign-flipping of AUC values per participant. We considered only FWE-corrected p-values below 0.05 (one-tailed) as significant.

A similar statistical testing procedure was applied to the temporal generalization analysis to test for temporal generalization effects trained on building/box at one time point and tested on scene/chair at another time point. Except for the AUC value of every sample coming from a time-by-time matrix, other steps in establishing clusters’ significance were the same as those in the category-level decoding mentioned above.

To further quantify the extent of off-diagonal generalization, we first defined a square window symmetric to the diagonal, covering exactly all the significant clusters, as a region of interest (ROI). Then, we compared the average AUC values in the upper triangular region within the ROI to those in the lower triangular region using a permutation-based paired-samples t-statistic.

To further localize the off-diagonal asymmetry, for every sample within the ROI [time * (time – 1)/2], we compared the AUC value in the upper triangular region (time i,j) with the corresponding AUC value in the lower triangular region (time j,i) using a paired-samples t-statistic. Then, adjacent samples were clustered based on a t-value threshold corresponding to an uncorrected p-value of 0.05, two-tailed (given that the asymmetry could go in both directions). The resulting cluster(s)’ FWE-corrected p-values were computed using the Monte Carlo method involving 10,000 random sign-flipping of AUC upper versus lower triangular difference values per participant. We considered only FWE-corrected p-values below 0.05 (one-tailed) as significant.

### fMRI study

#### Participants

Similar to the EEG experiment, we aimed to include 34 participants. A total of 34 participants were recruited through SONA, a psychology research related participants pool based at Radboud University, in exchange for monetary compensation. Four participants had to be excluded due to excessive head movement (head movement larger than 0.5 mm in both translational directions for five or more runs), resulting in a sample size of 30 participants (17 females, mean age = 24 ± 3.5 years). All participants reported having normal or corrected-to-normal vision and no history of neurological or psychiatric disorders. Written informed consent was obtained from all participants before the experiment. The experiment was approved by the local ethics committee (CMO Arnhem-Nijmegen).

#### MRI Acquisition

Anatomical and functional images were acquired using a 3T Siemens PrismaFit system (Siemens, Erlangen, Germany) equipped with a 32-channel headcoil. The entire MRI session lasted between 1.5 to 2.0 h, during which we acquired: (i) a T1-weighted anatomical scan (1 mm isotropic), (ii) one whole-brain functional localizer run to localize the PPA (3.5 mm isotropic; TR=1,500 ms). This run was used for an online analysis at the MRI scanner console to aid in slice positioning for the ultra-fast functional runs, (iii) one ultra-fast functional localizer run (2 mm isotropic; TR=140 ms), in which functional slices were positioned based on the online analysis of the whole-brain functional localizer (buildings vs. tools). This run served to independently localize the PPA from the slice positioning procedure and had the same scanning parameters as the following experimental runs, (iv) Six to seven ultra-fast functional runs to measure BOLD activity for different object categories (buildings, boxes, scenes, and chairs) during the experimental paradigm. In addition to the above runs, fieldmap images and one whole-brain volume were also acquired to correct for distortions and facilitate functional-anatomical image co-registration.

Anatomical images were acquired using a T1-weighted magnetization prepared rapid gradient echo (MP-RAGE) sequence (TR/echo time (TE) = 2300/3.03 ms, voxel size 1×1×1 mm, 8° flip angle). BOLD activity for the whole-brain functional localizer run was measured using T2*-weighted gradient echo planar imaging sequence (TR/TE = 1500/30 ms, 21 transversal slices, FOV 224 mm, voxel size 3.5×3.5×3.5 mm, 90° flip angle; slice gap = 0%). BOLD activity for the ultra-fast runs was measured using a T2*-weighted gradient echo planar imaging sequence (multiband acceleration factor = 3; TR/TE = 140/28 ms, 6 transversal slices, FOV 201 mm, voxel size 2.5×2.5×3 mm, 22° flip angle; slice gap = 20%). Notably, the fast TR was made possible by combining multiband imaging with a significant reduction in the number of slices.

#### Stimuli

For the functional localizer runs, stimuli consisted of colored images taken from 3 basic-level categories (scenes, tools, and chairs). Each category contained 20 exemplars. The colored images were resized and cropped to maintain a consistent retinal size. Specifically, images of tools and chairs were fit into a fixed-size squared mask and cropped; images of scenes were cropped by an aperture with the same diameter size as images of tools and chairs. The images were then superimposed on a phase-scrambled background. During the experiment, images with the phase-scrambled background were centrally displayed on a uniform gray background. The phase-scrambled background extended approximately 15 degrees and objects (and scenes) extended approximately 6 degrees.

For the experimental runs, stimuli were the same as those used in the EEG experiment (Figure 1A). Notably, none of the scene images in the functional localization runs were included in the experimental runs. All images were processed and displayed similarly to those in the functional localizer runs. Furthermore, there was one additional phase-scrambled image without any object superimposed, used as a blank trial to model baseline brain activity. Images of tools, instruments and hands were included for another experiment with a different purpose.

#### Experimental Design

The experiment consisted of three parts. The first part aimed to compute an online t-contrast (scenes > tools) at the MR console to aid the slice positioning of the subsequent ultra-fast scanning (Figure 3A). Participants were shown images of scenes and tools in a blocked design. Each block consisted of 20 images from the same category (scenes or tools). Each image was displayed for 450 ms, followed by a 300 ms interstimulus interval. Within the 20 images in a block, 2 images were immediate repetitions, evenly distributed across the image sequence. Participants were asked to detect the targets (i.e., repetitions) by pressing a button. The blocks of scenes and tools were displayed in a fixed alternating order, with the block of scenes always being the first block. There were 40 blocks in total. The actual blocks used to compute the online t-contrast varied across participants, ranging from 16 blocks (4 mins) to 40 blocks (10 mins), based on the discernibility of individual participants’ PPA activity (most participants’ PPA could be reliably discerned with 16 blocks of stimulation). After the PPA was reliably discerned in each participant, six transversal slices were positioned ventrally in the medial temporal lobe around the parahippocampal gyrus and adjusted to cover as much as possible of the bilateral PPA activation from the online t-contrast map. The slice positioning procedure took about 5 minutes.

Second, after positioning the slices in the bilateral PPA, participants underwent another functional localizer run with the ultra-fast fMRI sequence. The purpose of this functional localizer run was to independently localize the PPA for the following ultra-fast experimental runs. Specifically, the functional localizer run was similar to the first part, except that two additional conditions were added (chairs and fixation). It consisted of 24 blocks (six blocks for each of the four stimulus categories), totaling 6 minutes and 24 seconds (Figure 3A, left).

Third, participants completed 6 to 7 runs with the same ultra-fast fMRI sequence as in part 2. In each run, they were shown images of buildings, boxes, scenes, chairs, tools, instruments, and hands, as well as a blank image with only the phase-scrambled background. Each image was displayed for 300 ms, and the interstimulus interval (ISI) varied exponentially from 1000 to 4000 ms (mean ISI = 1700 ms). All 8 exemplar images from each of the 7 categories, along with the blank image (which was repeated 8 times), were displayed twice in each run, totaling 128 image displays per run. The order of images was fully randomized. Participants were asked to pay close attention to the images. After every 64 images, an additional probe stimulus was shown, and participants had to decide whether the probe stimulus had appeared in the immediately preceding 64 images by pressing yes/no buttons. They had a maximum of 5 seconds to make a response; any response beyond this limit was considered incorrect. A feedback screen indicating the correctness of their response appeared immediately after they made the response or exceeded the response limitation. There were 2 probes in each run. This recognition task was used to ensure participants stayed attentive, and performance data was not analyzed.

**Figure 3.**
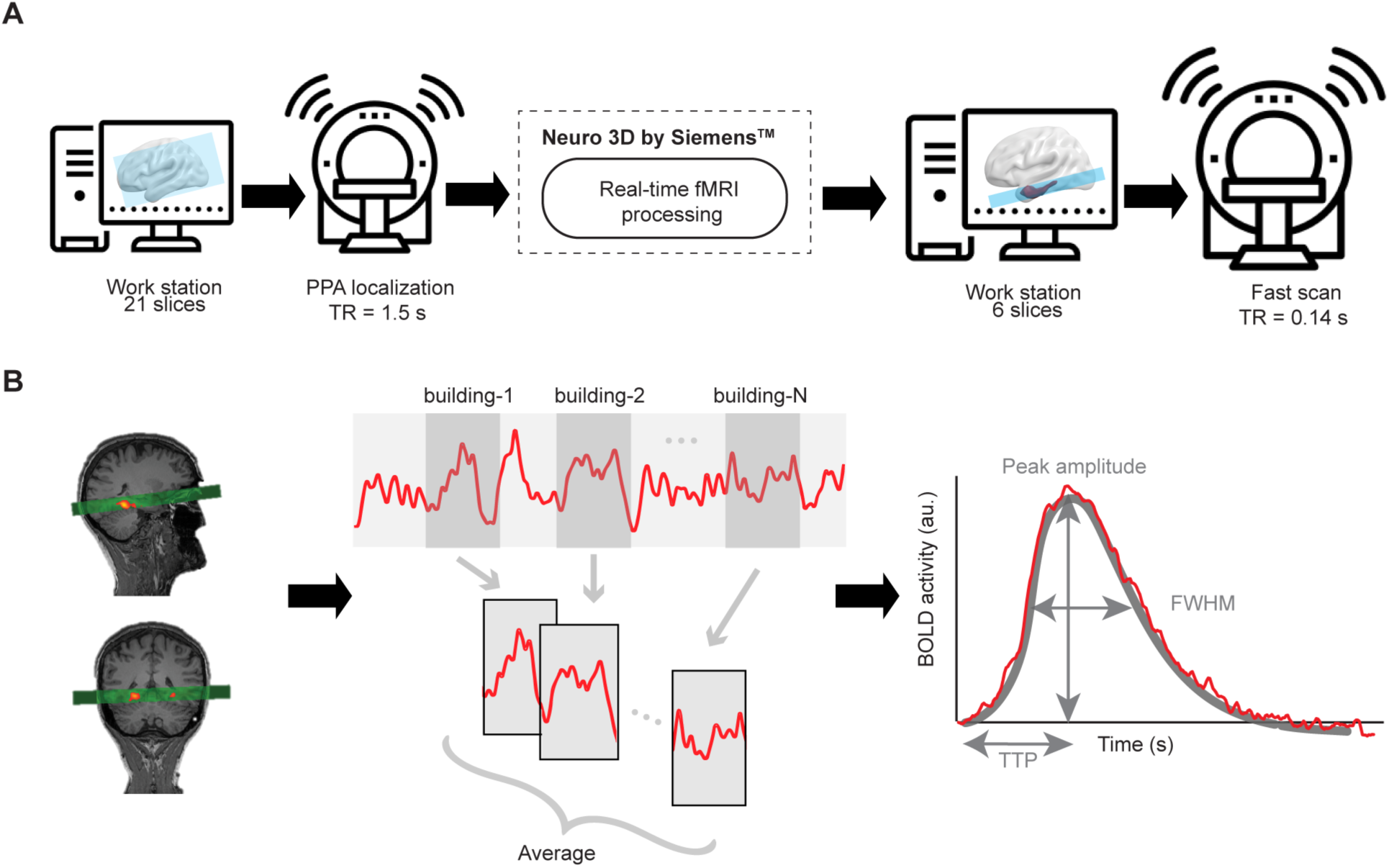
Experimental procedure and analysis outline of the ultra-fast fMRI experiment. (A) Schematic plot of the experimental procedure of ultra-fast fMRI. (B) Analysis outline of the ultra-fast fMRI. Left: One example participant’s PPA activation (red to yellow) in the functional localizer scan in the sagittal view (left) and coronal view (middle). Middle: the BOLD signal from the PPA was segmented and averaged across trials. Right: Schematic depiction of the procedure of HRF parameters estimation. The red line represents a hypothetical HRF time course, whereas the grey solid line represents the fitted single gamma function, with peak amplitude, TTP and FWHM as free parameters.

#### fMRI Preprocessing

Images from the ultra-fast sequence (including functional localizer and experimental runs) were preprocessed using a multiple-stage approach, considering the low signal-to-noise ratio (SNR) and unfavorable sampling of periodic physiological noise (e.g., respiratory and cardiac activity) inherent in fast fMRI (Chen et al., 2019). First, images were minimally preprocessed using FSL (Oxford, UK) including removing the first 28 volumes (∼ 4 s) to avoid magnetic field instability, motion correction (six-parameter affine transform, with the initial SBRef image of each run produced in multiband sequence as the reference image, given its higher image quality), and B0 distortion correction with fieldmap. For the functional localizer, an additional 5 mm Gaussian kernel spatial smoothing was applied. No spatial smoothing was performed for the experimental runs.

In the second stage of preprocessing, images from each experimental run were realigned to the SBRef image of the functional localizer run. GLMdenoise was then applied specifically for denoising the event-related experimental runs by estimating the noise regressors directly from the data (Kay et al., 2013). The resulting denoised 4D BOLD time series (indicated as denoiseddatà in the output of GLMdenoise toolbox) with the estimated noise contribution removed went into downstream analyses. However, for the functional localizer images, this method was inapplicable, so this step was omitted. Next, images from both the functional localizer run and experimental runs were bandpass filtered (0.01 – 0.15 Hz) to remove periodic physiological noise above 0.5 Hz.

In the last stage of preprocessing, independent component analysis (ICA) was performed for each run, which includes both functional localizer and experimental runs. Independent components (ICs) representing head movement were identified and removed through visual inspection by the experimenter, following the approach by Griffanti et al. (2017). All subsequent analyses were carried out in the native space of the images.

#### PPA Definition

A method similar to the Group-Constrained Subject-Specific (GSS) approach was employed to define each participant’s bilateral parahippocampal place area (PPA) as functional regions of interest (ROIs) (Julian et al., 2012). The GSS method helps define ROIs in individuals while minimizing subjectivity typically present in traditional handpicked ROI approaches. To do this, onsets and durations of different categorical blocks in the ultra-fast functional localizer run were convolved with a single-gamma hemodynamic response function (HRF) and fitted using a general linear model. For each participant, a one-sided t-contrast was calculated, comparing scene blocks to the average of tools and chairs blocks. The resulting statistical map was thresholded at p < 0.05 (one-tailed), uncorrected.

Next, the PPA probabilistic activation map (PAM) from the probabilistic functional atlas (Zhen et al., 2015) was thresholded at 50%, serving as an a priori constraint mask for the PPA. Individual PPAs were defined as the 30 highest scene-selective voxels after intersecting the a priori mask with each participant’s thresholded activation map. To assess the robustness of the results across different ROI sizes, the results were assessed at different ROI sizes from 10 to 70 voxels in increments of 10 voxels.

#### Amplitude, Time-to-Peak latency, and Full Width at Half Maximum

The time courses of the experimental runs were retrieved from the PPA ROIs and transformed into percent signal change unit by subtracting and dividing the time-averaged signal of each voxel. The time courses were then averaged across the voxels of the ROI for each participant and segmented into epochs spanning from 4 seconds before the onset of each stimulus to 15 seconds after. Epochs corresponding to the same stimulus condition were averaged across runs for each participant and baseline-corrected by subtracting the average of the pre-stimulus 4-second interval. Finally, the activity at corresponding time points in the blank background condition was subtracted from the epochs.

Given the intricate nature of neurovascular coupling, relying solely on a single parameter like Time-to-Peak latency (TTP) for interpretation can lead to oversimplification. Indeed, past studies have shown significant inter- and intra-subject correlations between TTP and the Full Width at Half Maximum (FWHM) (de Zwart et al., 2005; Siero et al., 2011). This suggests that these two parameters capture different, but interconnected, aspects of the temporal dynamics of the hemodynamic response. Therefore, to better encapsulate the multifaceted nature of temporal dynamics of hemodynamic responses, both TTP and FWHM, as well as fMRI response amplitude were set as free parameters and estimated by fitting a conventional single-gamma HRF function to the PPA time course, separately for each condition. To increase the signal-to-noise ratio, the time courses were averaged across participants and the group-averaged time courses were used for the fitting (all goodness-of-fit > 90%). The fitting was done using the curve_fit function as implemented in SciPy 0.18 (Ekman et al., 2017). The objective function was the sum of squared errors between the predicted and observed response. The TTP was constrained to a range between 3.8 and 8.6 seconds, which was 3 standard deviations around the group-averaged TTP based on a previous systematic study (Taylor et al., 2022). Similarly, the FWHM was constrained to a range between 1.8 and 6 seconds, 3 standard deviations around the group-averaged FWHM from the same study (Taylor et al., 2022).

#### Statistical Testing

To assess the statistical significance of amplitude, TTP and FWHM differences between various conditions, we used a permutation-based paired sample test. This was performed by first randomly permuting condition assignments of two individual-averaged time courses in each participant. Next, time courses were re-averaged across participants for each newly formed condition and fitted by the single-gamma HRF function to estimate amplitude, TTP and FWHM in these new conditions. The differences for amplitude, TTP and FWHM between the two conditions was recorded. This process was repeated 10,000 times, and a parameter (amplitude, TTP or FWHM) distribution was constructed under the null hypothesis that there was no significant difference between the two conditions. With this null distribution, we were able to assess the significance of difference between the two conditions of interest.

## Results

We conducted three studies involving independent groups of participants: a behavioral study measuring the perceptual similarity of a stimulus set consisting of buildings and boxes, an EEG decoding study, and an ultra-fast fMRI study (TR=140 ms).

### Behavioral study

We created a stimulus set consisting of eight pairs of visually matched buildings and boxes (Figure 1A). To confirm that buildings and boxes were indeed perceptually matched, across pairs, we conducted a behavioral study (N = 40 participants). In this study, participants were asked to indicate the location (left or right) of an “oddball” target among an array of spatially arranged distractors. These distractors were the same object shown in different sizes. Following previous research (Arun, 2012), 1/RT was used as a measure of perceptual dissimilarity between the oddball target and distractor objects. The oddball target and distractor can be any two different objects, such as visually matched buildings and boxes (matched objects, Figure 1B, left), visually unmatched buildings and boxes (unmatched objects, Figure 1B, middle), or objects of the same category (either buildings or boxes, Figure 1B, right).

After estimating the pairwise perceptual dissimilarity (or distance) between all pairs (averaged across two symmetrical oddball target and distractor assignments, Figure 4A), we first wanted to confirm that visually matched buildings and boxes indeed exhibited lower perceptual dissimilarity compared to unmatched ones. By grouping object combinations into matched and unmatched objects, we found that the average perceptual dissimilarity for matched objects was significantly lower than that for unmatched objects (*p < .001*, two-tailed permutation test; Figure 4B), showing that the visual matching manipulation decreased the perceptual distance of matched objects. More importantly, the average perceptual dissimilarity between objects from the same category (e.g., building-building) was equivalent to the perceptual dissimilarity between objects from different categories (*p = .999*, two-tailed permutation test; Figure 2B), indicating equal perceptual dissimilarity between exemplars within a category and between categories. This was only possible when exemplars from buildings and boxes were perceptually matched, otherwise the perceptual dissimilarity of objects within the same category would be lower than that between categories.

**Figure 4.**
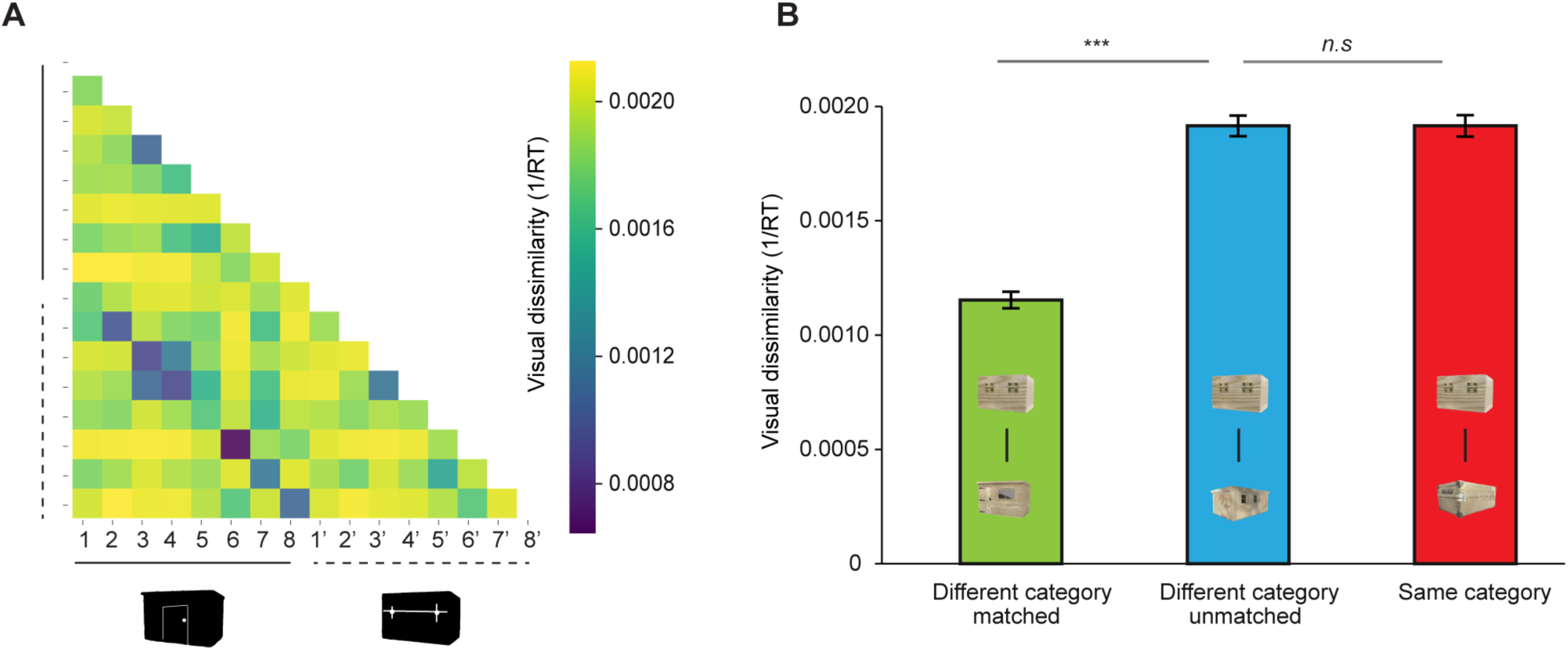
Pairwise perceptual dissimilarity matrix and averaged perceptual dissimilarity in each group. (A) Pairwise perceptual dissimilarity. Each element in the matrix represents a dissimilarity score between two different objects. (B) Averaged perceptual dissimilarity for matched different-category objects, unmatched different-category objects, and same-category objects. *** represents *p < .001*, n.s: non-significant. Error bars represent standard error.

### EEG study

After confirming that buildings and boxes were perceptually matched, next, capitalizing on EEG’s high temporal resolution, we investigated whether buildings evoke a selective response relative to boxes, and whether this selective response corresponds to scene-related processes. The same images of buildings and boxes used in the behavioral study were used in the EEG study. The images of buildings and boxes were further superimposed on a phase-scrambled background to equate image-level visual properties. Examination of image-level visual property metrics and activations from the layers of AlexNet confirmed that visual features were well-matched between buildings and boxes (Figures S1 and S2). Those images of buildings and boxes, together with scenes and chairs, were presented in random order while EEG signal was recorded (N = 32 participants). To make sure that participants stayed attentive during the experiment, they were asked to make an old/new judgement after every 6 images (Figure 2). EEG activity from posterior channels was used as input for linear classifiers trained at each time point. The classifier’s decoding performance was evaluated to indicate the presence of distinguishable information between two conditions.

#### Early and late distinction between scenes and chairs

We first examined when images of scenes and chairs elicited distinct EEG response patterns by evaluating the classification performance for scenes vs. chairs at each time point. As expected, based on previous M/EEG decoding studies (Cichy and Pantazis, 2017; Khaligh-Razavi et al., 2018), scenes and chairs were distinguishable from as early as 120 ms post-stimulus onset (*p < .001*, one-tailed, Figure 5, left). The distinction between scenes and chairs remained significant until the end of the analyzed time window (800 ms). This decoding likely reflects a mixture of both lower-level and higher-level differences between scenes and chairs.

**Figure 5.**
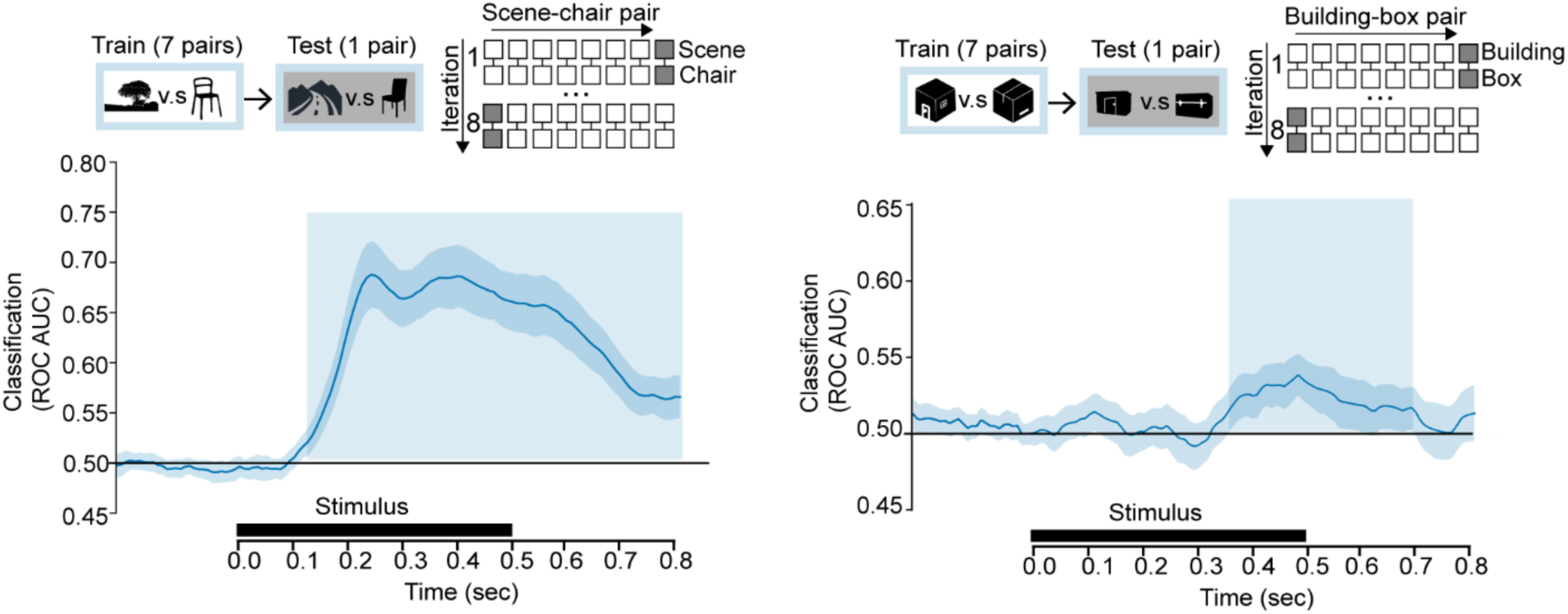
EEG experiment procedures and leave-one-pair-out EEG decoding results. Left: Scene vs. chair decoding. Right: Building vs. box decoding. The horizontal axis underneath represents post-stimulus onset time, and the black rectangle represents the duration of a stimulus. Y-axis indicates classification performance evaluated with ROC AUC. The dark blue area indicates ± 1 standard deviation. The light blue shaded area indicates time windows where classification performance is significantly above chance level.

#### Late distinction between buildings and boxes

Next, we examined whether EEG patterns could also distinguish between the closely matched buildings and boxes. Classification was done across pairs, such that the classifier was trained on 7 building-box pairs and tested on the held-out pair. In this way, the specific visual differences between individual buildings and boxes could not drive the decoding. Indeed, the behavioral study and image-level analyses (Figures S1, S2) showed that visual feature differences were not consistent across the pairs, such that they could not drive the cross-pair decoding. Significant decoding was observed relatively late in time, from 360 to 688 ms after stimulus onset (*p < .001*, one-tailed, Figure 5, right). We interpret this result as reflecting a higher-level cognitive process associated with large/stable (vs. small/manipulable) objects. Indeed, the relatively late decoding onset makes it unlikely that this result was driven by residual visual feature differences between buildings and boxes, which would be reflected in the first 250 ms of decoding (Carlson et al., 2013; Cichy and Pantazis, 2017; Proklova et al., 2019). These results were corroborated by qualitatively similar results from a representational similarity analysis in which perceptual similarity was regressed out (Figure S3).

#### Correspondence between buildings and scenes

To test whether the late building-specific activity patterns corresponded to scene-selective activity patterns, we examined whether the classifier trained on buildings vs. boxes generalized to distinguish scene vs. chairs. Results indicated that this was the case, despite the fact that these distinctions involved entirely different images and categories: the classifier trained on buildings vs. boxes could decode scenes vs. chairs from 336 to 680 ms after stimulus onset (*p < .001*, one-tailed, Figure 6A). To our knowledge, this provides the first EEG evidence for representational overlap between buildings and scenes.

#### Buildings indirectly activate scene-specific activity patterns

If the late processing for buildings (vs. boxes) reflected the indirect activation of scene-related processes (Mullally and Maguire, 2011), the building-specific pattern may correspond to a scene-specific pattern appearing earlier in time. To test this hypothesis, we performed a temporal generalization analysis, where building vs. box classifiers were trained on each time point and tested on each time point to distinguish scenes vs. chairs. Echoing the cross-decoding results, we observed a significant cluster in the temporal generalization matrix, demonstrating significant cross-decoding from buildings vs. boxes to scenes vs. chairs (*p = .002*, one-tailed, Figure 6B, left). Importantly, this cluster was not symmetrical with the diagonal, as shown by a significantly larger averaged AUC values in the upper triangular region than the lower triangular region within the square ROI covering the cluster (*p = .010*, one-tailed permutation test).

Further cluster-based analyses confirmed that the asymmetry was driven by a cluster peaking around 350 to 500 ms for the buildings (vs. boxes) and 200 to 300 ms for the scenes (vs. chairs; *p = .011*, one-tailed; Figure 6B, right). The centroid of the cluster indicates an average distinguishability timing of 398 ms for buildings vs. boxes. In contrast, the average timing for decoding scenes vs. chairs was earlier, at approximately 266 ms. This results in a temporal delay (ΔTime) of 132 ms.

**Figure 6.**
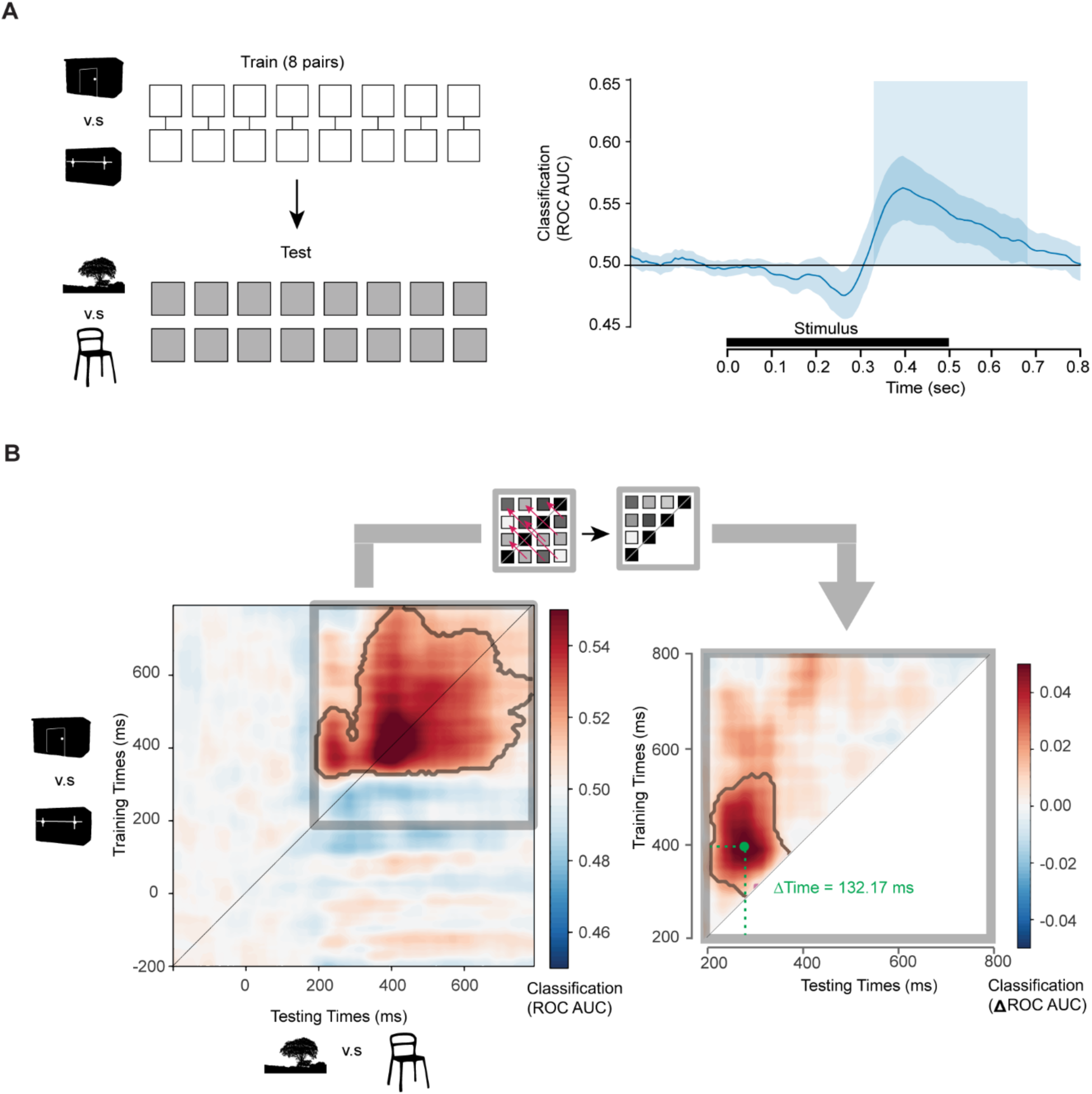
EEG cross-decoding results. (A) Cross decoding from building vs. box to scene vs. chair for the same time point. Left: Schematic plot of the cross-decoding training/test set arrangement. The 8 pairs of buildings and boxes were used as the training set to train a linear classifier, which was then tested on the 8 images of scenes and chairs. Right: Time course for buildings vs. boxes to scenes vs. chairs decoding. The horizontal axis underneath represents post-stimulus onset time, and the black rectangle represents the duration of a stimulus. Y-axis indicates classification performance evaluated with ROC AUC. The dark blue area indicates ± 1 standard deviation. The light blue shaded area indicates time windows where classification performance is significantly above chance level. (B) Temporal generalization analysis. Left: each point in the matrix represents the decoding performance of a classifier trained on buildings vs. boxes at the corresponding timepoint (y-coordinate) and tested on scenes vs. chairs at the corresponding timepoint (x-coordinate). The gray outline represents the contour of the cluster with significant above-chance decoding, whereas the gray rectangle represents the square covering the significant cluster symmetric to the diagonal. Right: Enlarged plot representing the point-by-point difference in decoding performance between upper-left triangle and lower-right triangle. The gray outline represents the contour of the cluster with a significant point-by-point difference in decoding.

In summary, we observed a distinguishable response to buildings relative to boxes in the EEG signal. The building-selective pattern emerged relatively late in time and corresponded to a scene-selective pattern that emerged earlier.

### fMRI study

While EEG offers high temporal resolution, its limited spatial resolution does not allow for determining the brain regions where buildings’ activation of scene-related processes occur. To investigate whether the scene-selective PPA responds selectively to buildings relative to the visually-matched boxes, and whether it shows a temporal delay for buildings, as observed in EEG, we next used fMRI (N = 30). Because of the relatively high spatial resolution of fMRI as compared with EEG, we could extract responses directly within the scene-selective PPA, rather than relying on scene-selective scalp-level patterns. We therefore analyzed the univariate response in the PPA rather than using decoding analysis. Indeed, the fMRI decoding equivalent of the EEG decoding analysis reported above would include voxels across the whole posterior brain (which was not covered by our ultra-fast scanning sequence). Also, in that case, decoding (e.g., scenes vs chairs) would primarily rely on the scene-selective peaks (i.e., PPA and other scene-selective regions) within this larger brain region. Another analysis option would be to decode categories within the PPA itself. However, this would not be comparable to the EEG analysis, as this decoding would be sensitive to fine-grained patterns within the PPA (which, for example, distinguish between individual scene categories; Walther et al., 2009) rather than the categorical distinctions (scenes, buildings) that were of interest here.

High temporal resolution was achieved mainly by reducing the number of scanning slices to only cover bilateral PPA. Briefly, the individual PPAs were first localized while participants were in the scanner (online localization). Then, the 6 slices were placed covering bilateral PPA (Figure 3A) to achieve a TR of 140 ms. With this high sampling rate, another functional localizer run was conducted (offline localization), followed by 6-7 main experiment runs. The offline localization was used to localize the PPA with the high sampling rate. The hemodynamic response of a specified category in the PPA was deconvolved by averaging trials corresponding to that category across all participants. We then fitted a single gamma function to the hemodynamic response to estimate the amplitude and temporal dynamic features including time-to-peak (TTP) latency and full-width-half-maximum (FWHM) for each stimulus category (Figure 3B). Note that we did not evaluate the latency of onset, another parameter often used to describe the temporal characteristics of the HRF. Previous studies have suggested that the latency of onset is impacted by amplitude (Thompson et al., 2014), whereas no correlations between TTP/FWHM and amplitude were observed (Miezin et al., 2000; Henson et al., 2002; Lindquist and Wager, 2007; Casanova et al., 2008; Thompson et al., 2014).

#### Higher PPA amplitude for buildings than boxes

While several studies have reported that the PPA responds more strongly to buildings (or large/stable objects more generally) than to other objects (Aguirre et al., 1998; Epstein and Kanwisher, 1998; Downing et al., 2006; Mullally and Maguire, 2011; Konkle and Oliva, 2012; Troiani et al., 2014), it is not known whether this selectivity is preserved when contrasting buildings with visually-matched controls. Because the buildings and boxes in the current study were matched on a range of low- and mid-level features (including rectilinearity), and EEG decoding showed no reliable differences in the first 300 ms of visual processing (indicating the absence of low- and mid-level visual feature differences that were consistent across exemplar pairs), the current fMRI experiment allowed us to address this question.

Results showed that buildings evoked a substantially stronger response than boxes in both the right PPA (PSC for buildings = .25%, PSC for boxes = .09%, *p < .001,* one-tailed permutation test, Figure 7A) and the left PPA (PSC for buildings = .22%, PSC for boxes = .09%, *p < .001*, one-tailed permutation test). Furthermore, this result was robust across ROI sizes (Figure S6). Overall, these results suggest that visual features do not fully explain the PPA response to buildings.

**Figure 7.**
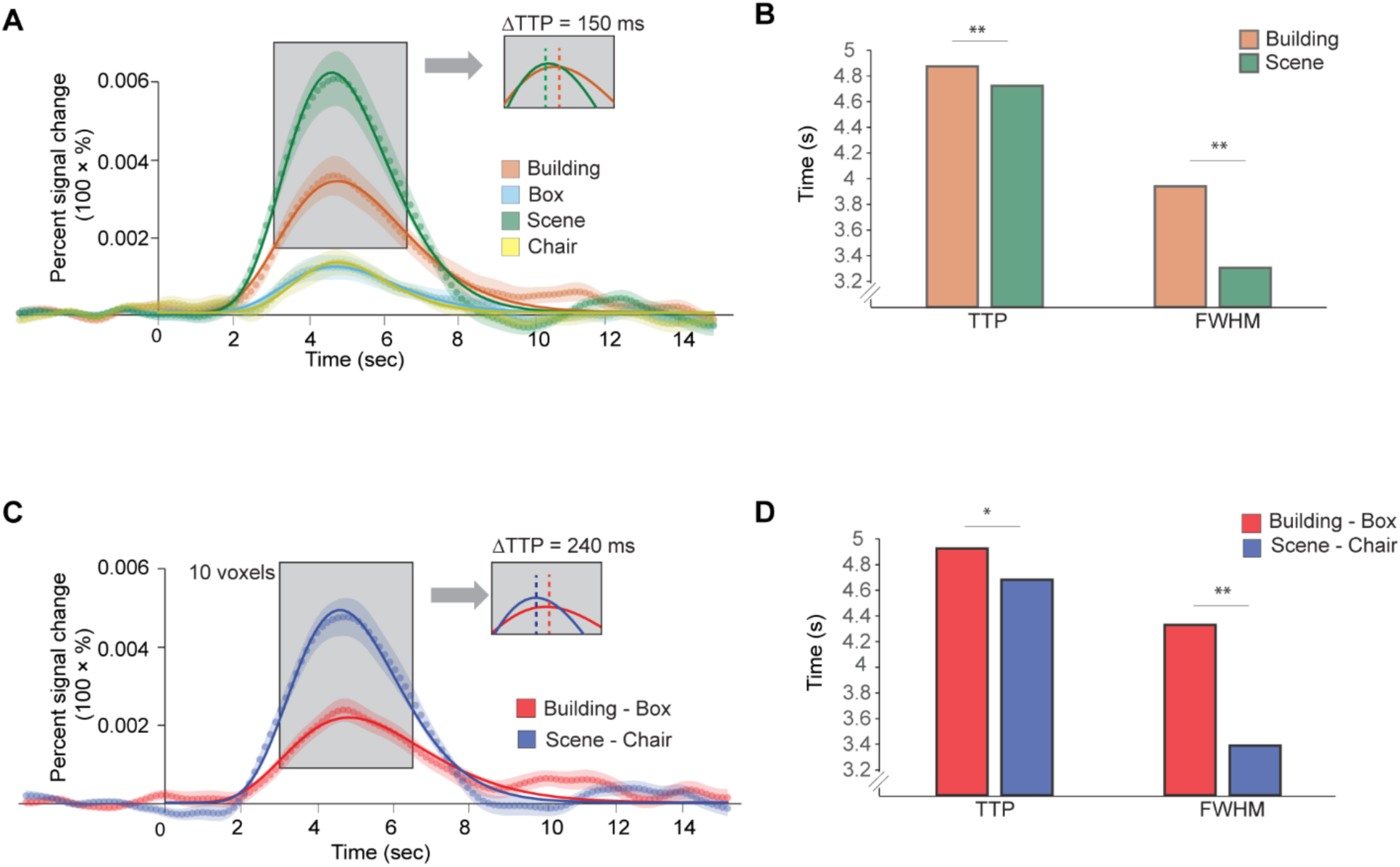
fMRI results. (A) HRF for buildings, boxes, scenes, and chairs for the right PPA. The dotted line represents the data in the corresponding condition, with the same-colored area representing ± 1 standard deviation. Solid lines represent the fitted HRFs (see Methods). The gray rectangle shows the peak latency difference between buildings and scenes. Note that in the gray inset, the fitted HRF for buildings and scenes were aligned regarding amplitude to illustrate the peak latency difference. (B) TTP and FWHM for buildings and scenes for the right PPA. (C) HRF for buildings minus boxes and scenes minus chairs for the right PPA. The dotted line represents the data in the corresponding condition, with the same-colored area representing ± 1 standard deviation. Solid lines represent the fitted HRFs (see Methods). The gray rectangle represents the peak latency difference between buildings and scenes. Note that in the gray inset, the fitted HRF for buildings and scenes were aligned regarding amplitude to illustrate the peak latency difference. (D) TTP and FWHM for buildings-boxes and scenes-chairs for the right PPA. *, ** represent *p < .05, 01*, respectively; between-subject error bars could not be plotted as significance was determined using permutation tests on group-averaged data.

#### Delayed and prolonged PPA activity for buildings relative to scenes

Next, we investigated whether the building-selective response, evidenced by the higher amplitude for buildings than boxes, was delayed and prolonged compared to the scene-selective response, as observed in the EEG study. To explore this, we first compared the temporal parameters of the fitted HRF functions for buildings and scenes separately (i.e., without contrasting responses with control categories). We expected that the response to buildings would show a later peak (reflected in TTP) and/or a longer duration (reflected in FWHM) than the response to boxes. The results confirmed this hypothesis. Evidenced by both TTP and FWHM, the fMRI responses to buildings were delayed (TTP for buildings = 4.88 s, TTP for scenes = 4.73 s, ΔTTP =150 ms, *p = .002*, one-tailed permutation test; Figure 7A, B) and prolonged (FWHM for buildings = 3.94 s, FWHM for scenes = 3.31 s, ΔFWHM = 630 ms, *p = .001*, one-tailed permutation test, Figure 7A, B) for the right PPA. A similar pattern was observed in the left PPA (TTP for buildings = 4.70 s, TTP for scenes = 4.58 s, ΔTTP =120 ms, *p = .009*, one-tailed permutation test; FWHM for buildings = 3.73 s, FWHM for scenes = 3.31 s, ΔFWHM = 420 ms, *p = .018*, one-tailed permutation test). Moreover, the results were consistent across ROI sizes (Figure S7).

To confirm these results, in a second analysis we compared the HRF parameters for the two contrasts (buildings-boxes, scenes-chairs). Also in this comparison, the response to buildings peaked later than the response to scenes in the right PPA (TTP for buildings = 4.92 s, TTP for scenes = 4.68 s, ΔTTP =240 ms, *p = .016*, one-tailed permutation test, Figure 7C, D). The response to buildings was also more prolonged than the response to scenes in the right PPA (FWHM for buildings = 4.33 s, FWHM for scenes = 3.39 s, ΔFWHM = 940 ms, *p = .002*, one-tailed permutation test, Figure 7C, D). For the left PPA, although numerically consistent, the differences were not significant (TTP for buildings = 4.70 s, TTP for scenes = 4.65 s, ΔTTP = 50 ms, *p = .34*, one-tailed permutation test; FWHM for buildings = 3.72 s, FWHM for scenes = 3.67 s, ΔFWHM = 50 ms, *p = .46*, one-tailed permutation test). Finally, the above results were consistent across ROI sizes (Figure S8). Together, these fMRI results are consistent with the EEG results: buildings (vs. boxes) activate the scene-selective PPA and this activation is delayed and/or prolonged relative to the scene-selective response itself.

## Discussion

Here, we used EEG and fMRI to measure a selective response to buildings relative to visually matched control stimuli (boxes). This response emerged relatively late during object processing and corresponded to the scene-selective response.

The current results inform our understanding of the representation of real-world object size in visual cortex. Previous studies showed that medial VTC, including PPA, responds selectively to large and stable objects such as buildings (Aguirre et al., 1998; Epstein and Kanwisher, 1998; Downing et al., 2006; Mullally and Maguire, 2011; Konkle and Oliva, 2012; Troiani et al., 2014). Similarly, using M/EEG, studies showed that real-world size could be decoded from around 150 ms after stimulus onset (Khaligh-Razavi et al., 2018; Wang et al., 2022). There is now convincing evidence that this selective response partly reflects the processing of low-or mid-level visual features that are typical of large objects (Andrews et al., 2010; Rajimehr et al., 2011; Nasr and Tootell, 2012; Nasr et al., 2014; Coggan et al., 2016; Long et al., 2018; Wang et al., 2022). To our knowledge, the current study is the first to show large-object (building) selectivity that is unlikely to reflect such visual feature differences. While individual stimuli obviously varied in terms of their visual features, buildings and boxes were matched in terms of image-level metrics across the stimulus set (e.g., retinal size, luminance, spatial frequency, rectilinearity, and AlexNet representations). Thus, there were no consistent features (e.g., rectilinearity) that were uniquely and consistently associated with either building or box categories. Furthermore, EEG decoding was done across exemplars, and building-selective EEG response patterns generalized to an entirely different image set consisting of scenes vs chairs. A categorical (building versus box) response was also observed when regressing out item-wise perceptual similarity from the behavioral visual search task (Figure S3). Finally, the finding that buildings and boxes evoked indistinguishable responses in the first 300 ms after stimulus onset confirms that the response was unlikely to be driven by systematic visual feature differences between buildings and boxes. Altogether, these results demonstrate that low-or mid-level features do not fully explain selectivity for buildings, suggesting an alternative route driving selectivity in PPA. This is in line with previous studies showing that the PPA response is modulated by the navigational relevance of objects in newly learned environments (Janzen and van Turennout, 2004; Schinazi and Epstein, 2010), and the finding of a selective PPA response to large objects in congenitally blind participants (He et al., 2013).

If not visual features, what could the selective response to buildings reflect? When considering this question, we need to account for the overlap between the response to buildings and scenes, as well as the temporal delay we observed for buildings relative to scenes. One possibility is that buildings evoke a sense of surrounding space (Mullally and Maguire, 2011). Evidence for this hypothesis comes from a study showing that large/stable objects were consistently rated as evoking a sense of surrounding space, and that such “space-defining” objects activated the PPA, even when they were merely imagined (Mullally and Maguire, 2011). This account readily explains the overlap between responses to buildings and scenes, as well as the temporal delay for buildings, since spatial information is more directly available from pictures of scenes. More generally, the current results raise the possibility that the PPA may not be directly involved in the recognition of buildings or large objects (based on visual feature processing), but rather reflects spatial information that is evoked by objects with certain characteristics (e.g., being large and stable); notably, these characteristics make these objects suitable as landmarks (Troiani et al., 2014), in line with the proposed role of the PPA in navigation (Epstein and Baker, 2019). We propose that objects (both small and large) are identified through a common object recognition pathway (e.g., the lateral occipital cortex), with contextual and/or spatial expectations activating PPA indirectly following processing in this pathway, i.e., after objects have been identified.

To optimize our chances of finding a strong PPA response, we chose objects (buildings and boxes) that differed both in size and in stability, based on previous studies showing that large/stable objects most strongly activate the PPA (Troiani et al., 2014) and evoke a sense of space (Mullally and Maguire, 2011). However, it is possible that one of these properties is sufficient to selectively activate the PPA (Hagen et al., 2026). Furthermore, our study does not rule out the possibility that other object properties also activate the PPA. For example, previous studies reported PPA activity for objects that are strongly linked to a specific scene context (Bar and Aminoff, 2003) and for objects that had been presented at decision points during the navigation of new environments (Janzen and van Turennout, 2004; Schinazi and Epstein, 2010). Finally, any object may activate the PPA when it serves to disambiguate scene background (Brandman and Peelen, 2019, 2023).

The current results are qualitatively in line with those of an intracranial EEG recording study of PPA in a small sample of patients (Bastin et al., 2013) Although responses were much earlier than observed in our study, similar to the current results, building-selective responses (∼170 ms) occurred later than scene-selective responses (∼80 ms). However, as the buildings and non-building objects were not matched in terms of their visual features, it is possible that the still relatively early response to buildings reflected selectivity to such visual features (Nasr et al., 2014; Long et al., 2018). Alternatively, building-selective responses could have reflected a mixture of bottom-up visual feature selectivity and top-down scene associations. Indeed, in our daily-life environments, conceptual object properties (e.g., real-world size, animacy) systematically covary with mid-level visual features (Wang et al., 2022), such that these features provide a useful shortcut for activating high-level object representations. By equating visual features across object conditions, this shortcut was not available in the current experiment, allowing us to isolate top-down input to the PPA and show that buildings can indirectly activate scene representations.

While our ultra-fast fMRI sequence only covered a narrow slice centered on the PPA, two additional scene-selective regions have been identified in visual cortex: the occipital place area (OPA) and the retrosplenial complex (RSC). These regions are strongly connected to each other and to the PPA, often showing similar functional profiles (Epstein & Baker, 2019), though there are also differences between these regions in terms of what scene properties they represent (Epstein & Baker, 2019; Dilks et al., 2022). Interestingly, like the PPA, large objects and buildings activate the OPA and RSC (Hagen et al., 2026; Konkle & Oliva, 2012), raising the question of whether delayed feature-independent building selectivity can also be observed in these regions. If so, these regions would also have contributed to the scalp-level EEG results observed here.

While here we investigated selectivity for buildings in medial VTC, we expect that the finding of delayed internally-driven category selectivity may similarly hold for other categories. For example, previous studies have provided evidence for selective responses to tools and body parts when contrasted with visually matched controls (Bracci and Beeck, 2016; Kaiser et al., 2016), and congenitally blind individuals show category-selective VTC responses to these same categories (Peelen et al., 2013; Kitada et al., 2014; Striem-Amit and Amedi, 2014). Therefore, the current results support the idea that category selectivity in visual cortex can be driven not only bottom-up, through the feedforward processing of visual features, but also top-down, likely reflecting input from connected downstream regions involved in functions such as tool use, reading, navigation, or social cognition (Martin, 2007; Mahon and Caramazza, 2011; Price and Devlin, 2011; Peelen and Downing, 2017). Similar to how the PPA may not be involved in recognizing or identifying large objects, other category-selective regions may also not encode the identity of objects of their preferred category. For example, tool-selective responses may not encode tool identity, but rather the hand actions associated with tools (Perini et al., 2014). Future studies could extend the current approach to other categories, including categories (e.g., faces) that appear to be primarily visually driven (Wang et al., 2015; Bi et al., 2016).

In summary, the current study provides evidence for a selective neural response to large objects (buildings) that is unlikely to reflect visual feature processing. The building-selective response consisted of a delayed activation of scene-selective mechanisms. These findings inform debates about the function and representational format of the PPA. More generally, they demonstrate that category selectivity in visual cortex can be dissociated from visual feature processing, with activity potentially reflecting feedback from domain-specific networks, following object recognition.

## Data & Code availability

The stimuli and data that support the findings of this study are available at: https://figshare.com/s/c08a2703a11125945b86. The code for reproducing the EEG and fMRI analyses can be accessed at: https://github.com/Yuan-fang/obj_timing_codes

## Author contributions

Y.Z. and M.P. designed research, Y.Z. and S.H. performed research, Y.Z. analyzed data, Y.Z. and M.P. interpreted the data, Y.Z. and M.P. wrote the paper.

## Acknowledgements

We would like to thank Dr. M. Ekman for suggestions regarding the fMRI data analysis, Dr. J.P. Marques for help with preparing the ultra-fast fMRI sequence, and P. Gaalman for help with data acquisition. We also thank Dr. B. Inglis at University of California, Berkeley for his generous help in solving an fMRI co-registration issue. This project received funding from the European Research Council (ERC) under the European Union’s Horizon 2020 research and innovation program (grant agreement No. 725970) awarded to M.V.P.. M.V.P. received support from the Netherlands Organization for Scientific Research (NWO-VICI: 231-057; https://doi.org/10.61686/CIOZQ88536).

## References

Aguirre GK, Zarahn E, D’Esposito M (1998) An Area within Human Ventral Cortex Sensitive to “Building” Stimuli: Evidence and Implications. Neuron 21:373–383.

Andrews TJ, Clarke A, Pell P, Hartley T (2010) Selectivity for low-level features of objects in the human ventral stream. NeuroImage 49:703–711.

Arun SP (2012) Turning visual search time on its head. Vision Res 74:86–92.

Bae G-Y, Luck SJ (2018) Dissociable Decoding of Spatial Attention and Working Memory from EEG Oscillations and Sustained Potentials. J Neurosci 38:409– 422.

Bainbridge WA, Oliva A (2015) Interaction envelope: Local spatial representations of objects at all scales in scene-selective regions. NeuroImage 122:408–16.

Bar M, Aminoff E (2003) Cortical Analysis of Visual Context. Neuron 38:347–358.

Bastin J, Vidal JR, Bouvier S, Perrone-Bertolotti M, Bénis D, Kahane P, David O, Lachaux J-P, Epstein RA (2013) Temporal Components in the Parahippocampal Place Area Revealed by Human Intracerebral Recordings. J Neurosci 33:10123– 10131.

Bi Y, Wang X, Caramazza A (2016) Object Domain and Modality in the Ventral Visual Pathway. Trends Cogn Sci 20:282–290.

Boccia M, Sulpizio V, Bencivenga F, Guariglia C, Galati G (2021) Neural representations underlying mental imagery as unveiled by representation similarity analysis. Brain Struct Funct 226:1511–1531.

Bracci S, Beeck HO de (2016) Dissociations and Associations between Shape and Category Representations in the Two Visual Pathways. J Neurosci 36:432–444.

Brandman T, Peelen MV (2019) Signposts in the Fog: Objects Facilitate Scene Representations in Left Scene-selective Cortex. J Cogn Neurosci 31:390–400.

Brandman T, Peelen MV (2023) Objects sharpen visual scene representations: evidence from MEG decoding. Cereb Cortex 33:9524–9531.

Carlson T, Tovar DA, Alink A, Kriegeskorte N (2013) Representational dynamics of object vision: The first 1000 ms. J Vis 13:1.

Casanova R, Ryali S, Serences J, Yang L, Kraft R, Laurienti PJ, Maldjian JA (2008) The impact of temporal regularization on estimates of the BOLD hemodynamic response function: a comparative analysis. NeuroImage 40:1606–1618.

Chen JE, Polimeni JR, Bollmann S, Glover GH (2019) On the analysis of rapidly sampled fMRI data. NeuroImage 188:807–820.

Cichy RM, Pantazis D (2017) Multivariate pattern analysis of MEG and EEG: A comparison of representational structure in time and space. NeuroImage 158:441–454.

Coggan DD, Liu W, Baker DH, Andrews TJ (2016) Category-selective patterns of neural response in the ventral visual pathway in the absence of categorical information. NeuroImage 135:107–114.

Dal Ben R (2023) SHINE_color: Controlling low-level properties of colorful images. MethodsX 11:102377.

de Leeuw JR (2015) jsPsych: A JavaScript library for creating behavioral experiments in a Web browser. Behav Res Methods 47:1–12.

de Zwart JA, Silva AC, van Gelderen P, Kellman P, Fukunaga M, Chu R, Koretsky AP, Frank JA, Duyn JH (2005) Temporal dynamics of the BOLD fMRI impulse response. NeuroImage 24:667–677.

Dilks DD, Kamps FS, Persichetti AS (2022) Three cortical scene systems and their development. Trends Cogn Sci 26:117–27.

Downing PE, Chan AW-Y, Peelen MV, Dodds CM, Kanwisher N (2006) Domain Specificity in Visual Cortex. Cereb Cortex 16:1453–1461.

Ekman M, Kok P, de Lange FP (2017) Time-compressed preplay of anticipated events in human primary visual cortex. Nat Commun 8:15276.

Epstein R, Kanwisher N (1998) A cortical representation of the local visual environment. Nature 392:598–601.

Epstein RA, Baker CI (2019) Scene Perception in the Human Brain. Annu Rev Vis Sci 5:373–397.

Gramfort A, Luessi M, Larson E, Engemann DA, Strohmeier D, Brodbeck C, Goj R, Jas M, Brooks T, Parkkonen L, Hämäläinen M (2013) MEG and EEG data analysis with MNE-Python. Front Neurosci 7:267.

Griffanti L, Douaud G, Bijsterbosch J, Evangelisti S, Alfaro-Almagro F, Glasser MF, Duff EP, Fitzgibbon S, Westphal R, Carone D, Beckmann CF, Smith SM (2017) Hand classification of fMRI ICA noise components. NeuroImage 154:188–205.

Hagen S, Zhao Y, Op de Beeck H, Peelen MV (2026) Dissociating representations of object shape, real-world size, and mobility in human visual cortex. bioRxiv 2026–7.

Haxby JV, Gobbini MI, Furey ML, Ishai A, Schouten JL, Pietrini P (2001) Distributed and Overlapping Representations of Faces and Objects in Ventral Temporal Cortex. Science 293:2425–2430.

He C, Peelen MV, Han Z, Lin N, Caramazza A, Bi Y (2013) Selectivity for large nonmanipulable objects in scene-selective visual cortex does not require visual experience. NeuroImage 79:1–9.

Henderson MM, Tarr MJ, Wehbe L (2023) A Texture Statistics Encoding Model Reveals Hierarchical Feature Selectivity across Human Visual Cortex. J Neurosci 43:4144–4161.

Henson RNA, Price CJ, Rugg MD, Turner R, Friston KJ (2002) Detecting Latency Differences in Event-Related BOLD Responses: Application to Words versus Nonwords and Initial versus Repeated Face Presentations. NeuroImage 15:83– 97.

Ishai A, Ungerleider LG, Haxby JV (2000) Distributed Neural Systems for the Generation of Visual Images. Neuron 28:979–990.

Janzen G, van Turennout M (2004) Selective neural representation of objects relevant for navigation. Nat Neurosci 7:673–677.

Julian JB, Fedorenko E, Webster J, Kanwisher N (2012) An algorithmic method for functionally defining regions of interest in the ventral visual pathway. NeuroImage 60:2357–2364.

Kaiser D, Azzalini DC, Peelen MV (2016) Shape-independent object category responses revealed by MEG and fMRI decoding. J Neurophysiol 115:2246–2250.

Kay K, Rokem A, Winawer J, Dougherty R, Wandell B (2013) GLMdenoise: a fast, automated technique for denoising task-based fMRI data. Front Neurosci 7:247.

Khaligh-Razavi S-M, Cichy RM, Pantazis D, Oliva A (2018) Tracking the Spatiotemporal Neural Dynamics of Real-world Object Size and Animacy in the Human Brain. J Cogn Neurosci 30:1559–1576.

King J-R, Dehaene S (2014) Characterizing the dynamics of mental representations: the temporal generalization method. Trends Cogn Sci 18:203–210.

Kitada R, Yoshihara K, Sasaki AT, Hashiguchi M, Kochiyama T, Sadato N (2014) The Brain Network Underlying the Recognition of Hand Gestures in the Blind: The Supramodal Role of the Extrastriate Body Area. J Neurosci 34:10096–10108.

Konkle T, Oliva A (2012) A Real-World Size Organization of Object Responses in Occipitotemporal Cortex. Neuron 74:1114–1124.

Kriegeskorte N, Mur M, Bandettini PA (2008) Representational similarity analysis - connecting the branches of systems neuroscience. Front Syst Neurosci 2:4.

Krizhevsky A, Sutskever I, Hinton GE (2012) ImageNet classification with deep convolutional neural networks. In: Advances in Neural Information Processing Systems 25, pp 1097–1105.

Lindquist MA, Wager TD (2007) Validity and power in hemodynamic response modeling: A comparison study and a new approach. Hum Brain Mapp 28:764– 784.

Long B, Yu C-P, Konkle T (2018) Mid-level visual features underlie the high-level categorical organization of the ventral stream. Proc Natl Acad Sci 115:E9015–E9024.

Mahon BZ, Caramazza A (2011) What drives the organization of object knowledge in the brain? Trends Cogn Sci 15:97–103.

Maris E, Oostenveld R (2007) Nonparametric statistical testing of EEG- and MEG-data. J Neurosci Methods 164:177–190.

Martin A (2007) The Representation of Object Concepts in the Brain. Annu Rev Psychol 58:25–45.

Miezin FM, Maccotta L, Ollinger JM, Petersen SE, Buckner RL (2000) Characterizing the Hemodynamic Response: Effects of Presentation Rate, Sampling Procedure, and the Possibility of Ordering Brain Activity Based on Relative Timing. NeuroImage 11:735–759.

Mullally SL, Maguire EA (2011) A New Role for the Parahippocampal Cortex in Representing Space. J Neurosci 31:7441–7449.

Nasr S, Echavarria CE, Tootell RBH (2014) Thinking Outside the Box: Rectilinear Shapes Selectively Activate Scene-Selective Cortex. J Neurosci 34:6721–6735.

Nasr S, Tootell RBH (2012) A Cardinal Orientation Bias in Scene-Selective Visual Cortex. J Neurosci 32:14921–14926.

Op de Beeck HP, Haushofer J, Kanwisher NG (2008) Interpreting fMRI data: maps, modules and dimensions. Nat Rev Neurosci 9:123–135.

Peelen MV, Bracci S, Lu X, He C, Caramazza A, Bi Y (2013) Tool Selectivity in Left Occipitotemporal Cortex Develops without Vision. J Cogn Neurosci 25:1225– 1234.

Peelen MV, Downing PE (2017) Category selectivity in human visual cortex: Beyond visual object recognition. Neuropsychologia 105:177–183.

Perini F, Caramazza A, Peelen MV (2014) Left occipitotemporal cortex contributes to the discrimination of tool-associated hand actions: fMRI and TMS evidence. Front Hum Neurosci 8:591.

Portilla J, Simoncelli EP (2000) A Parametric Texture Model Based on Joint Statistics of Complex Wavelet Coefficients. Int J Comput Vis 40:49–70.

Price CJ, Devlin JT (2011) The Interactive Account of ventral occipitotemporal contributions to reading. Trends Cogn Sci 15:246–253.

Proklova D, Kaiser D, Peelen MV (2019) MEG sensor patterns reflect perceptual but not categorical similarity of animate and inanimate objects. NeuroImage 193:167– 177.

Rajimehr R, Devaney KJ, Bilenko NY, Young JC, Tootell RBH (2011) The “Parahippocampal Place Area” Responds Preferentially to High Spatial Frequencies in Humans and Monkeys. PLOS Biol 9:e1000608.

Ritchie JB, Wardle SG, Vaziri-Pashkam M, Kravitz DJ, Baker CI (2026) Rethinking category-selectivity in human visual cortex. Cogn Neurosci 17:49–76.

Romei V, Brodbeck V, Michel C, Amedi A, Pascual-Leone A, Thut G (2008) Spontaneous Fluctuations in Posterior α-Band EEG Activity Reflect Variability in Excitability of Human Visual Areas. Cereb Cortex 18:2010–2018.

Schinazi VR, Epstein RA (2010) Neural correlates of real-world route learning. NeuroImage 53:725–735.

Siero JC, Petridou N, Hoogduin H, Luijten PR, Ramsey NF (2011) Cortical Depth-Dependent Temporal Dynamics of the BOLD Response in the Human Brain. J Cereb Blood Flow Metab 31:1999–2008.

Striem-Amit E, Amedi A (2014) Visual Cortex Extrastriate Body-Selective Area Activation in Congenitally Blind People “Seeing” by Using Sounds. Curr Biol 24:687–692.

Taylor AJ, Kim JH, Ress D (2022) Temporal stability of the hemodynamic response function across the majority of human cerebral cortex. Hum Brain Mapp 43:4924–4942.

Thompson SK, Engel SA, Olman CA (2014) Larger neural responses produce BOLD signals that begin earlier in time. Front Neurosci 8:159.

Troiani V, Stigliani A, Smith ME, Epstein RA (2014) Multiple Object Properties Drive Scene-Selective Regions. Cereb Cortex 24:883–897.

Walther DB, Caddigan E, Fei-Fei L, Beck DM (2009) Natural scene categories revealed in distributed patterns of activity in the human brain. J Neurosci 29:10573–81.

Wang R, Janini D, Konkle T (2022) Mid-level Feature Differences Support Early Animacy and Object Size Distinctions: Evidence from Electroencephalography Decoding. J Cogn Neurosci 34:1670–1680.

Wang X, Peelen MV, Han Z, He C, Caramazza A, Bi Y (2015) How Visual Is the Visual Cortex? Comparing Connectional and Functional Fingerprints between Congenitally Blind and Sighted Individuals. J Neurosci 35:12545–12559.

Zhen Z, Yang Z, Huang L, Kong X, Wang X, Dang X, Huang Y, Song Y, Liu J (2015) Quantifying interindividual variability and asymmetry of face-selective regions: A probabilistic functional atlas. NeuroImage 113:13–25.

