## Supplementary Material for "Turning an object into a scene: buildings activate scene-selective visual cortex independently of visual features"

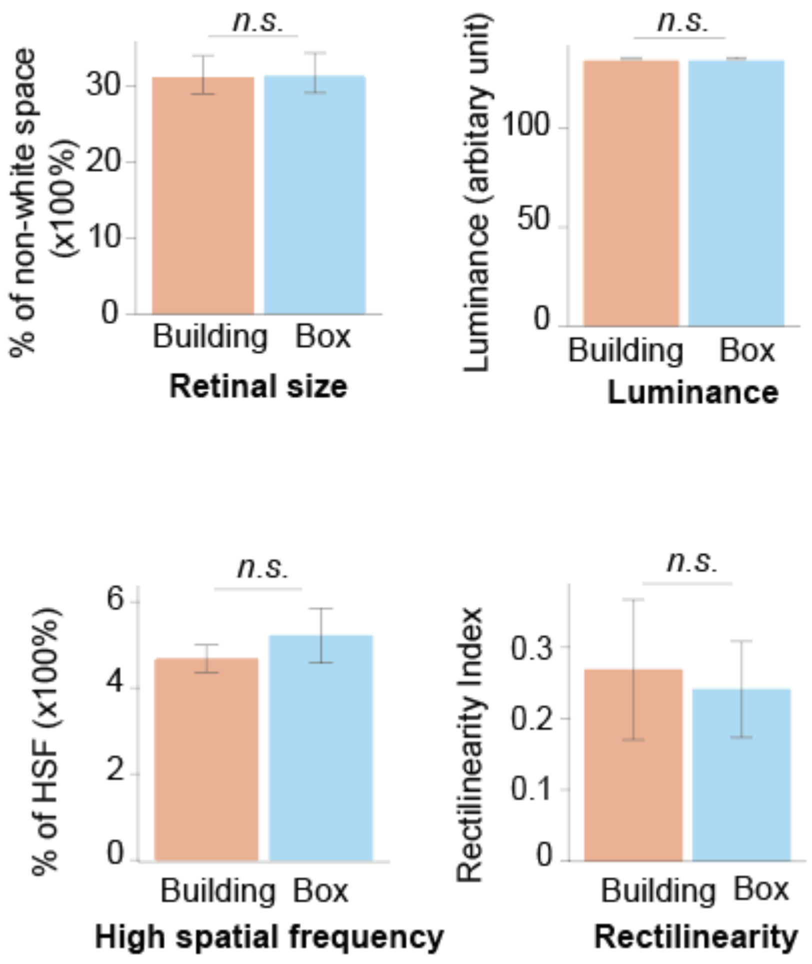


**Supplementary Figure 1.** Low/mid-level visual properties comparison between building and box images.

Upper left panel: building and box images have comparable retinal size ($p$ = .594), as indexed by % of non-white space. Upper right panel: building and box images have comparable luminance ($p$ = .852). Lower left panel: building and box images have comparable high spatial frequency (HSF, $p$ = .570), as indexed by % of HSF (defined as above 75 cycles per image). Lower right panel: building and box images have comparable rectilinearity index ($p$ = .828), computed based on Nasr et al., 2017 (https://github.com/cechava/Rectilinearity_Toolbox). Note that all the p-values are based on within-pair permutation-based two-tailed t-test (number of permutations = 10,000).

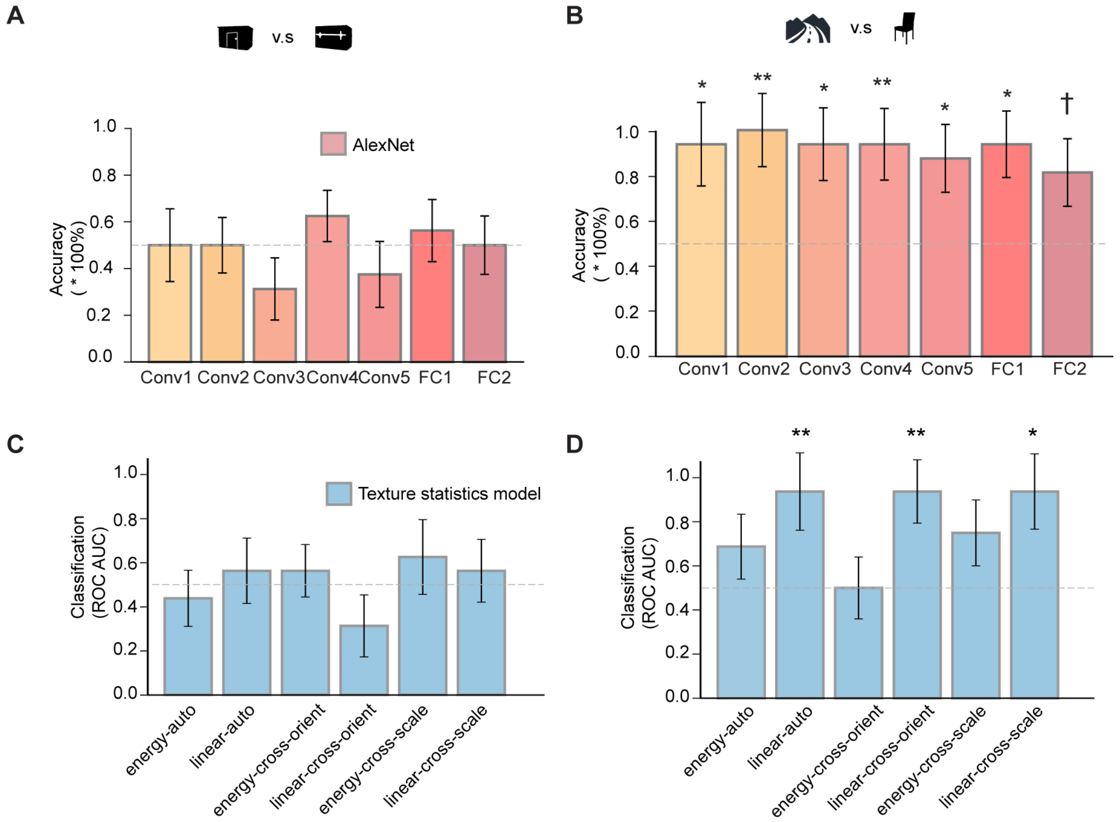


**Supplementary Figure 2.** (A, B) Decoding accuracy across all layers of AlexNet for building vs. box and scene vs. chair. (C, D) Results of the P-S model comparison.

(A) Decoding accuracy for buildings vs boxes. In each layer, one pair of building and box images was left out, an SVM classifier was trained on the remaining pairs and tested on the left-out pair. This process was iterated across all the 8 building-box image pairs. The accuracy was computed by averaging accuracies of the 8 iterations. The p-values were based on within-pair permutation-based classification accuracies (number of permutations = 1000, two-tailed). Conv1, $p = 1.0$; Conv2, $p = 1.0$; Conv3, $p = .261$; Conv4, $p = .432$; Conv5, $p = .495$; FC1, $p = .80$; FC2, $p = 1.0$

(B) Decoding accuracy for scenes vs chairs. Conv1, $p = .02$; Conv2, $p = .006$; Conv3, $p = .02$; Conv4, $p = .007$; Conv5, $p = .017$; FC1, $p = .01$; FC2, $p = .071$. The horizontal dashed line in both panels denotes chance-level decoding accuracy (50%).

(C) The P–S model, a texture statistics model constructed to extract multiple mid-level features with a steerable pyramid, provides the most comprehensive account of mid-level features (Portilla & Simoncelli, 2000). Importantly, the P–S model has proven useful in explaining fMRI responses to naturalistic stimuli in category-selective regions, including the PPA (Henderson et al., 2023). Moreover, the "texforms", a stimulus set preserving objects' mid-level features while being unrecognizable, were generated from mid-level visuo-statistical features of the P–S model and are widely used to probe mid-level features' unique contributions to category selectivity related to object size (Long et al., 2018; Wang et al., 2022). In lieu of the above, for each image, all the mid-level features specified in the P-S model were extracted within each hypothetical pRFs for the PPA, following a similar procedure as in Henderson et al., 2023. With those feature values pooled across all the pRFs, a leave-one-pair-out cross-validation scheme was applied to examine whether buildings and boxes could be reliably classified across all those mid-level features. As expected, buildings cannot be reliably distinguished from boxes above the chance level (i.e., classification accuracy: 50%) across all the mid-level features (*p*s > 0.2, nonparametric permutation-based test using N = 1000 permutations), further indicating that mid-level features were well matched between buildings and boxes.

(D) In contrast, scenes and chairs, distinct in many obvious visual features, can be successfully classified based on features including 'linear-auto' (classification accuracy: 94%, *p* = .008), 'linear-cross-orient' (classification accuracy: 94%, *p* = 0.009), and 'linear-cross-scale' (classification accuracy: 94%, *p* = 0.016).

Conv#: convolutional layer; FC#: fully connected layer. ** represents *p < .01*, * represents *p < .05*, † represents *p < .1*

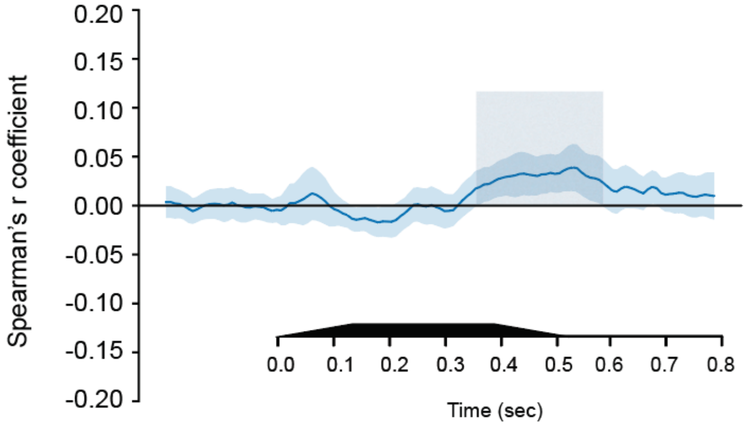


**Supplementary Figure 3.** EEG representational similarity analysis of categorical object representation after regressing out perceptual similarity for each time point.

At each time point, a partial Spearman's correlation coefficient was computed between the representational dissimilarity matrix (RDM) derived from EEG patterns across posterior channels for each object and the categorical relationship matrix, controlling for the perceptual similarity matrix, derived from the visual search task (1/RT). The light blue shaded area indicates time windows where correlation is significantly above 0. The horizontal axis underneath represents post-stimulus onset time, and the black rectangle represents the duration of a stimulus.

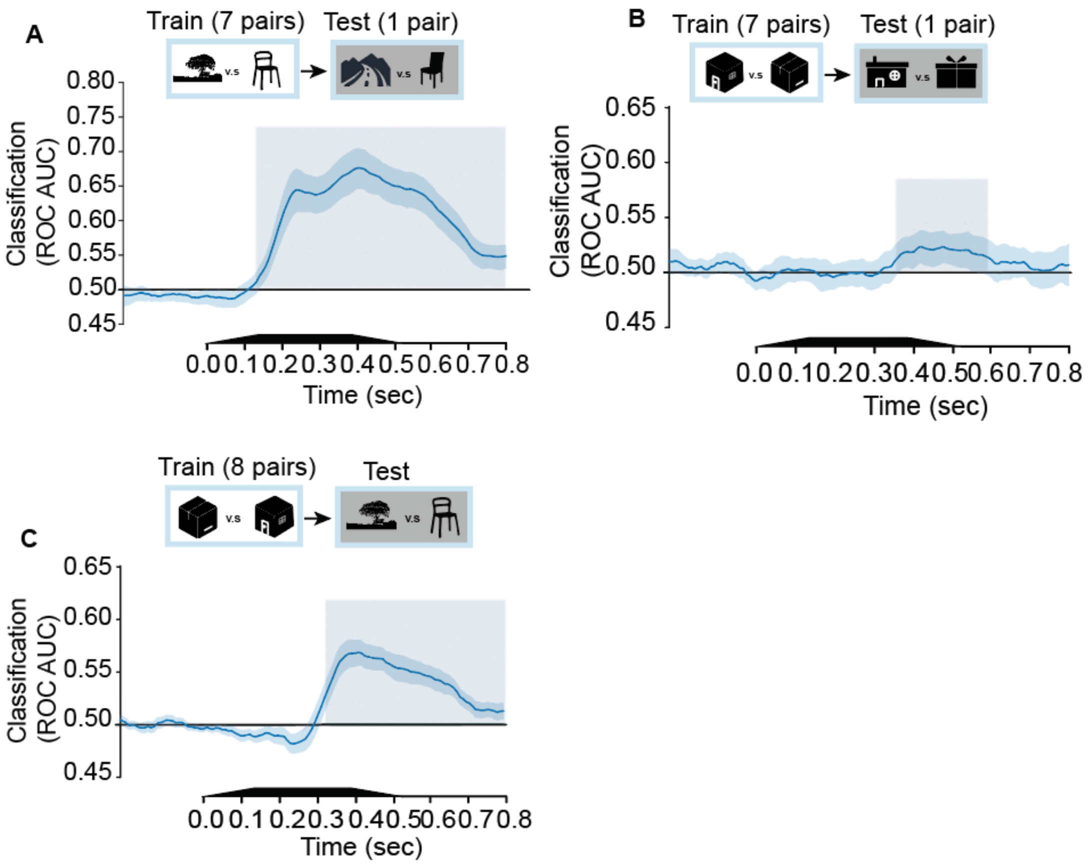


**Supplementary Figure 4.** EEG decoding results with all channels.

(A) Scene vs. chair decoding with all EEG channels. (B) Building vs. box decoding with all EEG channels. (C) Cross-decoding from building vs. box to scene vs. chair for corresponding time points. The horizontal axis underneath represents post-stimulus onset time, and the black rectangle represents the duration of a stimulus. Y-axis indicates classification performance evaluated with ROC AUC. The dark blue area indicates ± 1 standard deviation. The light blue shaded area indicates time windows where classification performance is significantly above chance level.

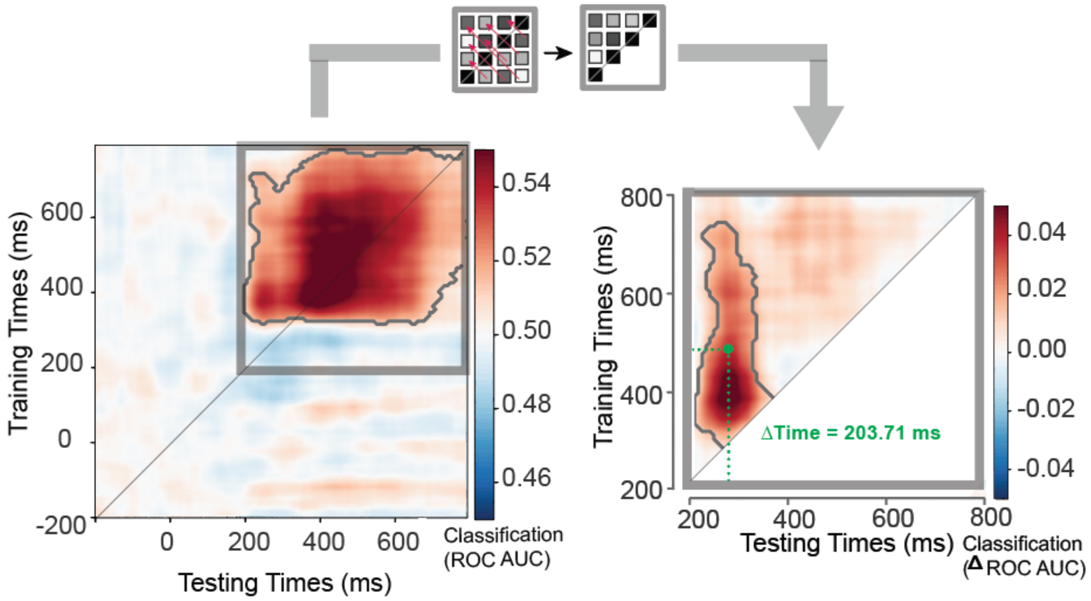


**Supplementary Figure 5.** EEG temporal generalization results with all channels.

Left: each point in the matrix represents the decoding performance of a classifier trained on buildings vs. boxes at one timepoint (y-coordinate) and tested on scenes vs. chairs at another timepoint (x-coordinate). The gray outline represents the contour of the cluster with significant above-chance decoding, whereas the gray rectangle represents the square covering the significant cluster symmetric to the diagonal. Right: Enlarged plot representing the point-by-point difference in decoding performance between upper-left triangle and lower-right triangle. The gray outline represents the contour of the cluster with a significant point-by-point difference in decoding.

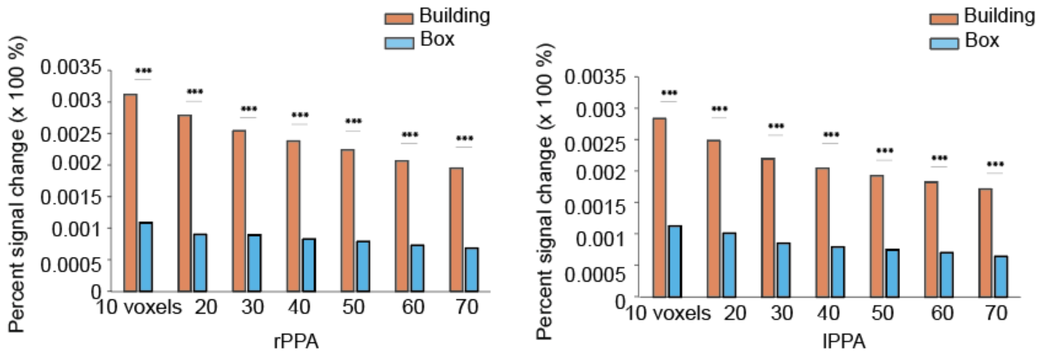


**Supplementary Figure 6.** fMRI response amplitude for buildings and boxes across different ROI sizes in the right and left PPA.

fMRI response amplitude was computed across different ROI sizes, ranging from 10 to 70 voxels in increments of 10 voxels. Voxel sizes were defined as the N highest scene-selective voxels after intersecting the a-priori PPA probabilistic activation mask with each participant’s thresholded scene-selective map. The resulting individual-averaged PPA time courses at different ROI sizes were further averaged across participants. The group-averaged time courses at different ROIs were used to estimate the amplitude by fitting a single gamma function. rPPA: right PPA; lPPA: left PPA. *** represents $p < .001$.

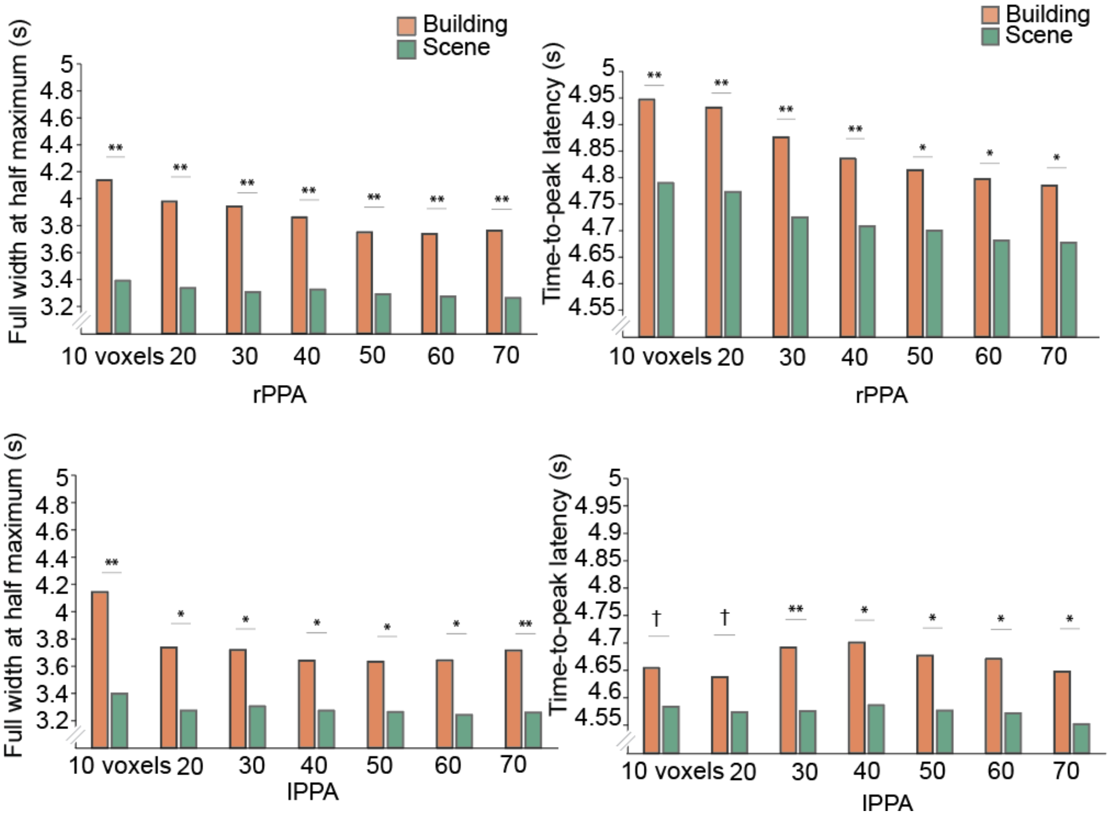


**Supplementary Figure 7.** FWHM and TTP for buildings and scenes across different ROI sizes in the right and left PPA.

FWHM and TTP were estimated across different ROI sizes, ranging from 10 to 70 voxels in increments of 10 voxels. Voxel sizes were defined as the N highest scene-selective voxels after intersecting the a-priori PPA probabilistic activation mask with each participant’s thresholded scene-selective map. The resulting individual-averaged PPA time courses at different ROI sizes were further averaged across participants. The group-averaged time courses at different ROIs were used to estimate the FWHM and TTP by fitting a single gamma function. rPPA: right PPA; lPPA: left PPA. ** represents $p < .01$, * represents $p < .05$,  and † represents $p < .1$.

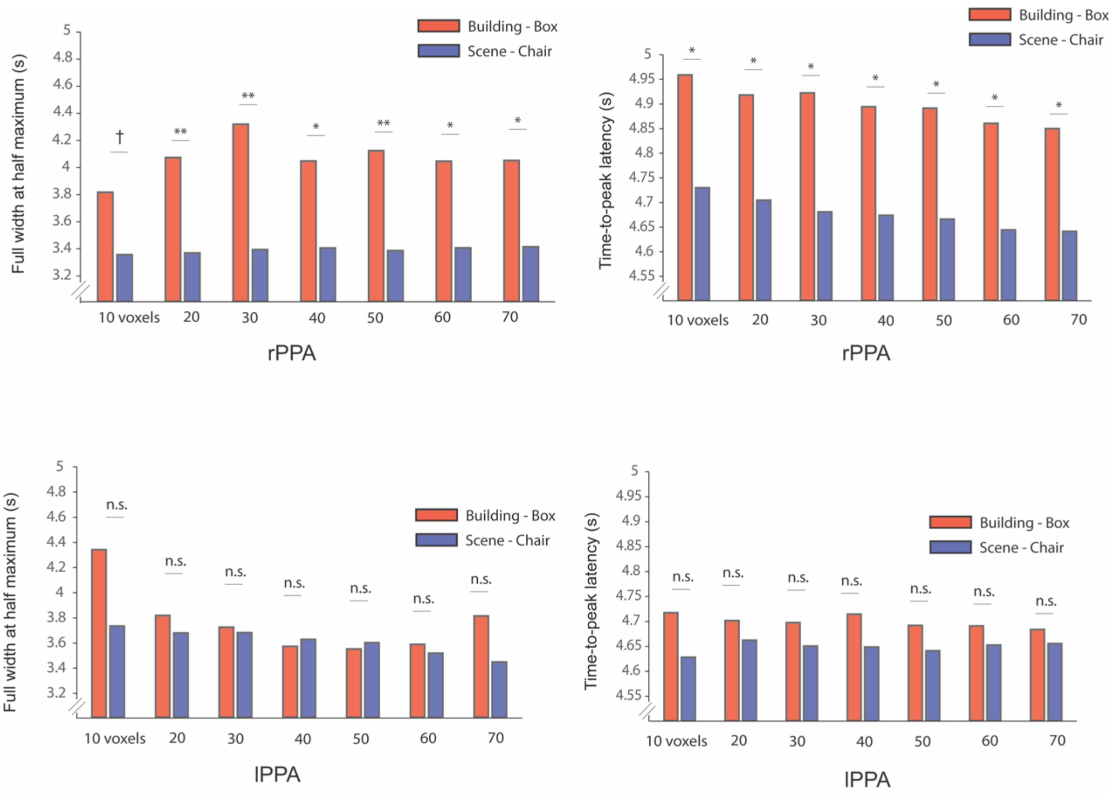


**Supplementary Figure 8.** FWHM and TTP for buildings vs. boxes and scenes vs. chairs across different ROI sizes in the right and left PPA.

FWHM and TTP were estimated across different ROI sizes, ranging from 10 to 70 voxels in increments of 10 voxels. Voxel sizes were defined as the N highest scene-selective voxels after intersecting the a-priori PPA probabilistic activation mask with each participant’s thresholded scene-selective map. The resulting individual-averaged PPA time courses at different ROI sizes were further averaged across participants. The group-averaged difference time courses between buildings vs. boxes or scenes vs chairs at different ROIs were used to estimate the FWHM and TTP by fitting a single gamma function. rPPA: right PPA; lPPA: left PPA. ** represents $p < .01$, * represents $p < .05$, † represents $p < .1$ , and n.s. represents $p > .1$.
